# Paleocene origin of a streamlined digestive symbiosis in leaf beetles

**DOI:** 10.1101/2023.12.12.571295

**Authors:** Marleny García-Lozano, Christine Henzler, Miguel Ángel González Porras, Inès Pons, Aileen Berasategui, Christa Lanz, Heike Budde, Kohei Oguchi, Yu Matsuura, Yannick Pauchet, Shana Goffredi, Takema Fukatsu, Donald Windsor, Hassan Salem

**Author notes:** Correspondence: Hassan Salem.

## Abstract

Timing the acquisition of a beneficial microbe relative to the evolutionary history of its host can shed light on the adaptive impact of a partnership. Here, we investigated the onset and molecular evolution of an obligate symbiosis between Cassidinae leaf beetles and *Candidatus* Stammera capleta, a γ-proteobacterium. Residing extracellularly within foregut symbiotic organs, *Stammera* upgrades the digestive physiology of its host by supplementing plant cell wall-degrading enzymes. We observe that *Stammera* is a shared symbiont across tortoise and hispine beetles that collectively comprise the Cassidinae subfamily, despite differences in their folivorous habits. In contrast to its transcriptional profile during vertical transmission, *Stammera* elevates the expression of genes encoding digestive enzymes while in the foregut symbiotic organs, matching the nutritional requirements of its host. Symbiont acquisition during the Paleocene (∼62 Mya) did not coincide with the origin of Cassidinae beetles, despite the widespread distribution of *Stammera* across the subfamily. Early-diverging lineages lack the symbiont and the specialized organs that house it. Reconstructing the ancestral state of host-beneficial factors revealed that *Stammera* encoded three digestive enzymes at the onset of symbiosis, including polygalacturonase – a pectinase that is universally shared. While non-symbiotic cassidines encode polygalacturonase endogenously, their repertoire of plant cell wall-degrading enzymes is more limited compared to symbiotic beetles supplemented with digestive enzymes from *Stammera*. Highlighting the potential impact of a symbiotic condition and an upgraded metabolic potential, *Stammera*-harboring beetles exploit a greater variety of plants and are more speciose compared to non-symbiotic members of the Cassidinae.

## Introduction

Folivores must contend with a diet rich in recalcitrant plant polymers such as cellulose and pectin ^1^. Tortoise beetles (Coleoptera: Chrysomelidae: Cassidinae) overcome these challenges by encoding cellulases endogenously ^2^, while outsourcing their pectinolytic metabolism to *Candidatus* Stammera capleta, a γ-proteobacterial symbiont ^3–5^.

Tortoise beetles house *Stammera* extracellularly in foregut symbiotic organs, in addition to ovary-associated glands to ensure transmission ^3,5,6^. Encoded within the symbiont’s drastically reduced genome (0.21 Mb) are plant cell wall-degrading enzymes that upgrade the digestive ability of the beetle host ^3,4,7^. Females vertically propagate the symbiosis by depositing a *Stammera*-bearing ‘caplet’ at the anterior pole of each egg ^3,5,8^. As caplet consumption initiates *Stammera* infection during embryo development ^9^, its experimental removal disrupts the symbiont’s transmission cycle, yielding aposymbiotic insects ^3,8^. Symbiont loss results in a diminished digestive capacity and low larval survivorship ^3,8^, highlighting the obligate dependence of tortoise beetles on *Stammera*.

The Cassidinae represents a highly diverse subfamily of leaf beetles, with more than 6000 herbivorous species occupying a variety of ecological guilds ^10^. The monophyletic group includes exophagous tortoise beetles, along with hispine beetles that are largely leaf-mining, and were formerly classified as a separate subfamily ^10–12^. A suite of morphological and behavioural traits differentiates both groups ^10^, but, critically, tortoise beetles vary from hispines in their host-plant use ^11–13^. Hispines feed predominantly on monocotyledonous plants such as grasses and palms, whereas tortoise beetles coevolved with a dicotyledonous flora ^10^, reflecting a derived dietary shift that corresponded with the emergence and diversification of a then-novel niche during the Cretaceous ^14,15^.

Previous efforts characterizing the Cassidinae-*Stammera* symbiosis focused exclusively on dicot-feeding taxa, suggesting that the partnership may have originated with, and is restricted to, tortoise beetles ^3,4,6^. Given their divergent nutritional ecology, it remained unclear whether tortoise and hispine beetles are both hosts to *Stammera* ^4,6,16^, despite their shared ancestry ^10^. Notably, if symbiont acquisition did not coincide with the origin of the subfamily, how do non-symbiotic cassidines process a leafy diet in the absence of *Stammera*-encoded pectinases?

Here, we determine the origin of the Cassidinae-*Stammera* symbiosis and emphasize the divergent strategies facilitating a folivorous lifestyle across a highly speciose clade of beetles. We do so by (i) reconstructing host-symbiont phylogenetic relationships spanning a wider and more representative distribution of Cassidinae tribes, including hispine lineages, (ii) assessing the *Stammera* pangenome across 50 strains to examine patterns of molecular evolution and testing for signatures of selection, (iii) reconstructing the ancestral configuration of host-beneficial factors, (iv) quantifying the symbiont’s transcriptional dynamics relative to beetle development and nutritional requirements, and (v) estimating the timing of *Stammera* acquisition and investigating its impact on the diversification of its herbivorous host.

## Results and discussion

### *Stammera* is a shared symbiont across tortoise and hispine beetles

Initial descriptions of the *Stammera*-Cassidinae symbiosis examined five tortoise beetle tribes, including the Cassidini, Mesomphaliini, Ischyrosonychini, Omocerini, and Notoscanthini ^3,4,6^. We extended our characterization of the partnership to 13 tribes (totaling 55 species) (Figure 1 A-C; Table S1), emphasizing hispine clades considered to represent early-diverging Cassidinae lineages, including the Alurnini and Spilophorini ^10^.

**Figure 1.**
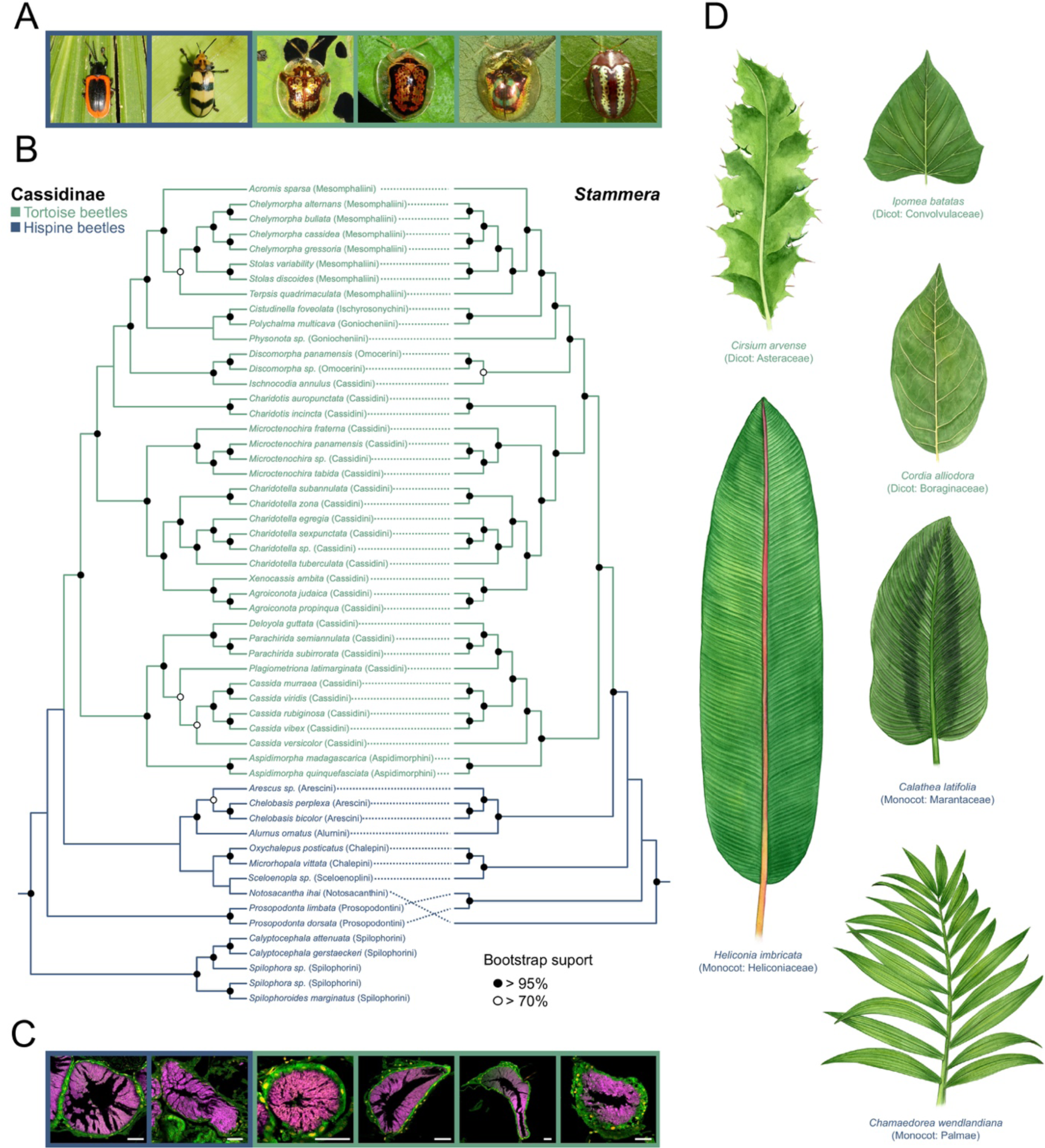
*Stammera* is a shared symbiont across tortoise and hispine beetles. (**A**) Hispine (dark blue outlines) and tortoise beetles (light blue outlines) from left to right: *Prosopodonta limbata, Chelobasis bicolor, Deloyala guttata*, *Microctenochira tabida*, *Charidotella zona* and *Chelymorpha alternans*. (**B**) Tanglegram depicting co-cladogenesis between Cassidinae beetles and their *Stammera* symbionts based on Maximum Likelihood (ML) phylogenies. Host tribes are indicated in parentheses. The Cassidinae phylogeny is based on a concatenated alignment of 15 mitochondrial genes, whereas the *Stammera* phylogeny was constructed used a concatenated alignment of 61 single-copy core genes. Node coloration reflects bootstrap support. Full detailed ML phylogenies including outgroups along with their complementary Bayesian phylogenetic trees for host and symbiont are included in Figures S4 and S3. (**C**) Fluorescence *in situ* hybridization (FISH) cross-sections of foregut symbiotic organs of the Cassidinae beetle species outlined above, targeting *Stammera* (magenta: 16 rRNA) and host (green: 18 rRNA) cells against the DAPI counterstain (yellow). Scale bars (50 μm) are included for reference. (**D**) Illustrations portraying representative host plants of hispine (Monocots: *Chamaedorea wendlandiana*, *Heliconia imbricata,* and *Calathea latifolia*) and tortoise beetles (Dicots: *Cordia alliodora*, *Cirsium arvense*, and *Ipomoea batatas*).

Metagenomic sequencing revealed the presence of *Stammera* in both tortoise and hispine beetles. Of the 55 Cassidinae species examined here, 50 are hosts to *Stammera* (Figure 1B). This is complemented by the presence of morphologically conserved symbiotic organs in hispines relative to the *Stammera*-bearing structures previously described in tortoise beetles (Figure 1C) ^3,5,6^. Both groups maintain *Stammera* extracellularly in specialized symbiotic organs connected to the foregut-midgut junction (Figure 1C). Reflecting the maternal inheritance of *Stammera* ^3,8^, we observe near-strict co-cladogenesis between the Cassidinae and their symbionts (Figure 1B) highlighted by 40 co-speciation events relative to 9 potential transfers across host lineages and 2 losses. This is consistent with insect symbionts that are transmitted vertically ^17^, as demonstrated in aphids ^18^, cicadas ^19^, stinkbugs ^20^, and bat flies ^21^.

While most hispine beetles surveyed in our study do harbor *Stammera* (Figure 1A-C), members of the most basal Spilophorini tribe lack the symbiont (Figure 1B) and the corresponding *Stammera*-bearing organs. Our findings indicate that symbiont acquisition was not synchronous with the origin of the host clade (Figure 1B). But while *Stammera* acquisition did not coincide with the origin of the Cassidinae, the digestive symbiosis predated the obligate monocot-to-dicot evolutionary transition that is concomitant with the rise and subsequent diversification of tortoise beetles (Figure 1B).

A comparative analysis spanning 50 *Stammera* genomes revealed that their sizes are drastically and consistently reduced, ranging from 216 to 340 Kb, and bearing only 201-317 protein-coding genes. Both parameters are positively correlated (Spearman’s rank correlation, rho = 0.934, *p* < 0.001) (Figure S1) as observed in other bacterial genomes ^22^, including insect symbionts ^23^. We also note that *Stammera* associating with tortoise beetles possesses the smallest genomes, in contrast to symbionts characterized in hispines belonging to the Alurnini, Chalepini and Prosopodontini tribes (Table S2), which possess the largest. In addition to a chromosome, most *Stammera* strains carry 1 to 2 plasmids ranging in size between 2.5 to 9 Kb. While plasmids predominantly encode genes for the host-beneficial factors supplemented by *Stammera*, we observe the loss of these extrachromosomal elements in a subset of species (e.g. *Stammera* from *Terpsis quadrimaculatta*), followed by their integration into the symbiont chromosome.

To understand how *Stammera* metabolic potential compares across hispine and tortoise beetles, we examined shared orthologues and gene loss following a pangenome analysis as implemented in anvi’o ^24^. Our analysis revealed a pangenome composed of 503 gene clusters, of which 125 (24.9%) are shared by all *Stammera* strains (core genes), compared to the 295 (58.6%) clusters representing the accessory gene set along with 83 (16.5%) singletons (Dataset 1) (Figure 2). Categories related to basic informational processes (replication, transcription, and translation) and posttranslational modification are highly represented within the core genome, whereas gene clusters underlying energy production, metabolism, and cell wall biogenesis are prevalent in the accessory genome (Figure S2). Such variation is observed in *Stammera* with the smallest genomes, suggesting that gene loss, rather than acquisition and rearrangement, might drive these differences, as recombination is expected to be minimal in clonal populations of vertically-transmitted symbionts ^25^. While 315 orthologous genes are shared between *Stammera* from tortoise and hispine beetles, a large proportion of genes are also uniquely present in each group (71 and 117, respectively) (Figure 3A). Upon comparing the Clusters of Orthologous Groups (COG) annotations across both groups, we identified pathways that are statistically enriched in *Stammera* genomes from hispine beetles. These pathways included lipoate biosynthesis (q-value < 0.01), pyruvate oxidation (q-value<0.01), isoleucine, leucine, valine biosynthesis (q-value < 0 .05), and menaquinone biosynthesis (q-value < 0.05). In contrast, no pathways were statistically enriched in *Stammera* genomes from tortoise beetles, consistent with the observation that gene loss was unidirectional throughout the molecular evolution of the symbiont. Finally, by examining gene order across representative symbiont genomes spanning the Cassidinae, we note that *Stammera* chromosomes are highly syntenic (Figure 3B). The high level of synteny, coupled with the monophyly of *Stammera* within the Enterobacteriaceae (Figure 1B; Figure S3), point to a single origin of symbiosis with the Cassidinae.

**Figure 2.**
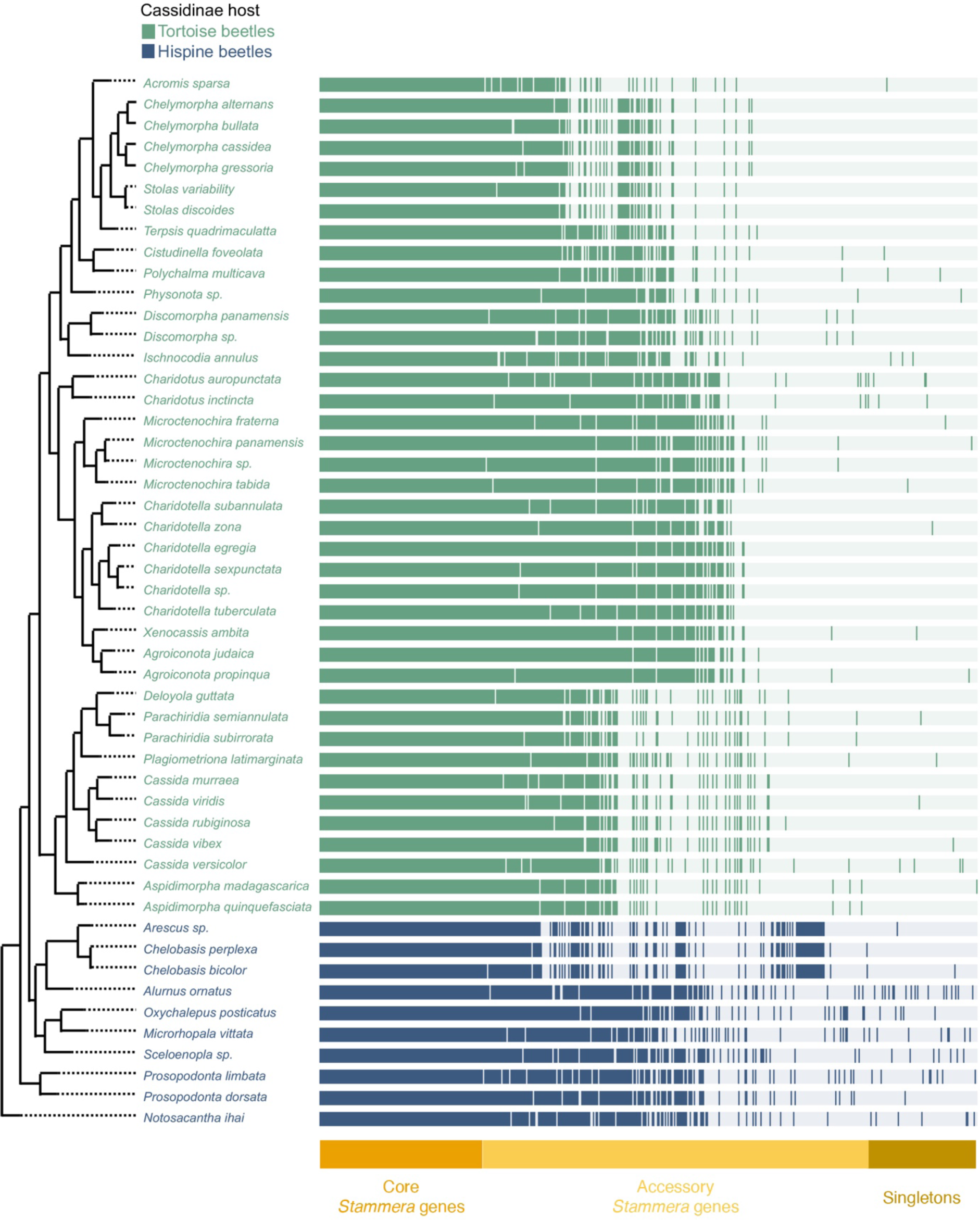
Core and accessory genes across the *Stammera* pangenome. Comparative genomics of 50 *Stammera* strains depicting the distribution of core, accessory and singleton genes. Each bar represents a *Stammera* genome from one Cassidinae host, and each layer illustrates the presence or absence of a gene cluster across the different genomes. Genomes are ordered according to a Maximum Likelihood *Stammera* phylogeny constructed using 124 single copy core genes. Core genes indicate gene clusters identified in all genomes, and accessory genes represent gene clusters discretely distributed but present in at least one genome. Singletons are genes present in just one genome.

**Figure 3.**
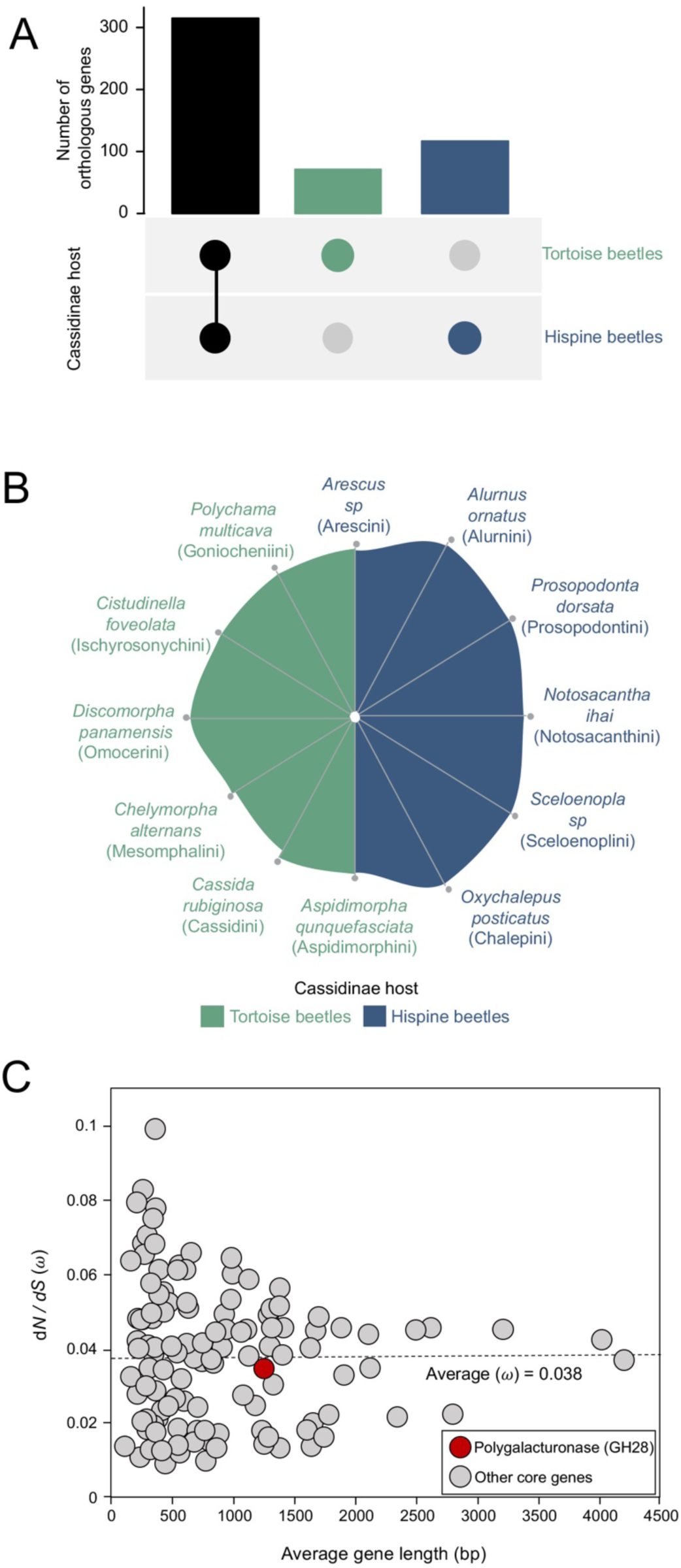
*Stammera* genomic features and molecular evolution. (**A**) UpSet plot of shared and non-shared gene content between *Stammera* from tortoise (light blue) and hispine beetles (dark blue). (**B**) Hive plot depicting gene order conservation spanning representative *Stammera* strains across the 12 Cassidinae host tribes surveyed in this study. (**C**) Relationship between average gene length and nonsynonymous (d*N*) to synonymous (d*S*) substitution values for core genes (gray) including polygalacturonase (red), a pectinase. Average gene length was calculated across all *Stammera* species. Dashed line represents the average rate (ω) of d*N*/d*S* substitutions across the orthologous symbiont genes. Abbreviation: GH, glycoside hydrolase.

### *Stammera* genes experiencing strong purifying selection

Genome-wide tests for selection can help identify genes underlying co-adaptation between a symbiont and its host ^18,26,27^. For example, by estimating the rate (ω) of nonsynonymous (d*N*) to synonymous (d*S*) substitutions acting on *Buchnera* genes, Chong and Moran ^18^ revealed a small subset of loci undergoing positive selection (ω > 1). These featured *Buchnera* membrane proteins that are highly expressed within the symbiotic organs of aphids ^28^, possibly facilitating interactions at the host-symbiont interface ^18^. In contrast, purifying selection (ω < 0.1) plays key role in preserving the functionality of long-term partnerships by purging deleterious mutations impacting critical functions, as highlighted for the obligate symbionts of leafhoppers ^26^, cicadas ^29^, and earthworms ^30^.

We leveraged the 50 *Stammera* genomes available in our study to examine the signatures of selection acting on shared loci by estimating ω across whole genes. Using the M0 model as implemented in PAML ^31^, this initial estimate revealed that *Stammera* genomes are experiencing strong purifying selection (average ω = 0.038, [range = 0.008-0.098, n = 124). In contrast, we observe no support for relaxed (0.95 < ω > 0.1) or positive selection (ω > 1). As specific codons related to intrinsic protein function may be experiencing different signatures relative to the whole gene, we additionally measured ω across codon sites and compared the site-based models M1a (nearly neutral) and M2a (positive selection). ω values were indicative of relaxed purifying selection (average ω = 0.119), revealing the absence of any positively selected sites within orthologous *Stammera* genes.

Strong selective constrains are described across several symbiont lineages, preserving key cellular functions, along with biosynthetic pathways that are essential for host development ^27^. Purifying selection is critical for the stability of obligately co-dependent partnerships and a driver of genome evolution across endosymbionts such as *Blattabacterium* in cockroaches ^32^ and *Thiodiazotropha* in clams ^33^, as well as beneficial extracellular microbes such as *Verminephrobacter* in earthworms ^30^. Gene categories underlying informational processing, chaperones, and host-beneficial factors typically exhibit the strongest levels of purifying selection ^34^. We observe that many of these genes are evolving under similar selective constraints in *Stammera* (Figure 3C; Dataset 2). Of the 20 *Stammera* genes exhibiting the lowest ω values (0.009-0.018), 17 are involved in transcription, translation and replication (e.g. *rho*, *infA* and *polA*), two underlie posttranslational modification (e.g. *smpB* and *rsmD*) and one is involved in amino acid transport and metabolism (e.g. *sufS*) (Dataset 2). Symbiont pathways encoding key metabolites for the host can experience similar selective pressures, including *Buchnera* genes involved in amino acid biosynthesis for aphids, or the nitrogen metabolism of *Blochmannia* in ants and *Wigglesworthia* in tsetse flies ^32^. We examined the signature of selection acting on polygalacturonase (Figure 3C) – a pectinase targeting nature’s most abundant pectic substrate, homogalacturonan – and the sole host-beneficial factor shared by all *Stammera* strains surveyed here (Figure 4). We observe that the pectinase is more constrained in its evolutionary change (ω = 0.035) than the average of all core genes (Figure 3C; Dataset 2).

**Figure 4.**
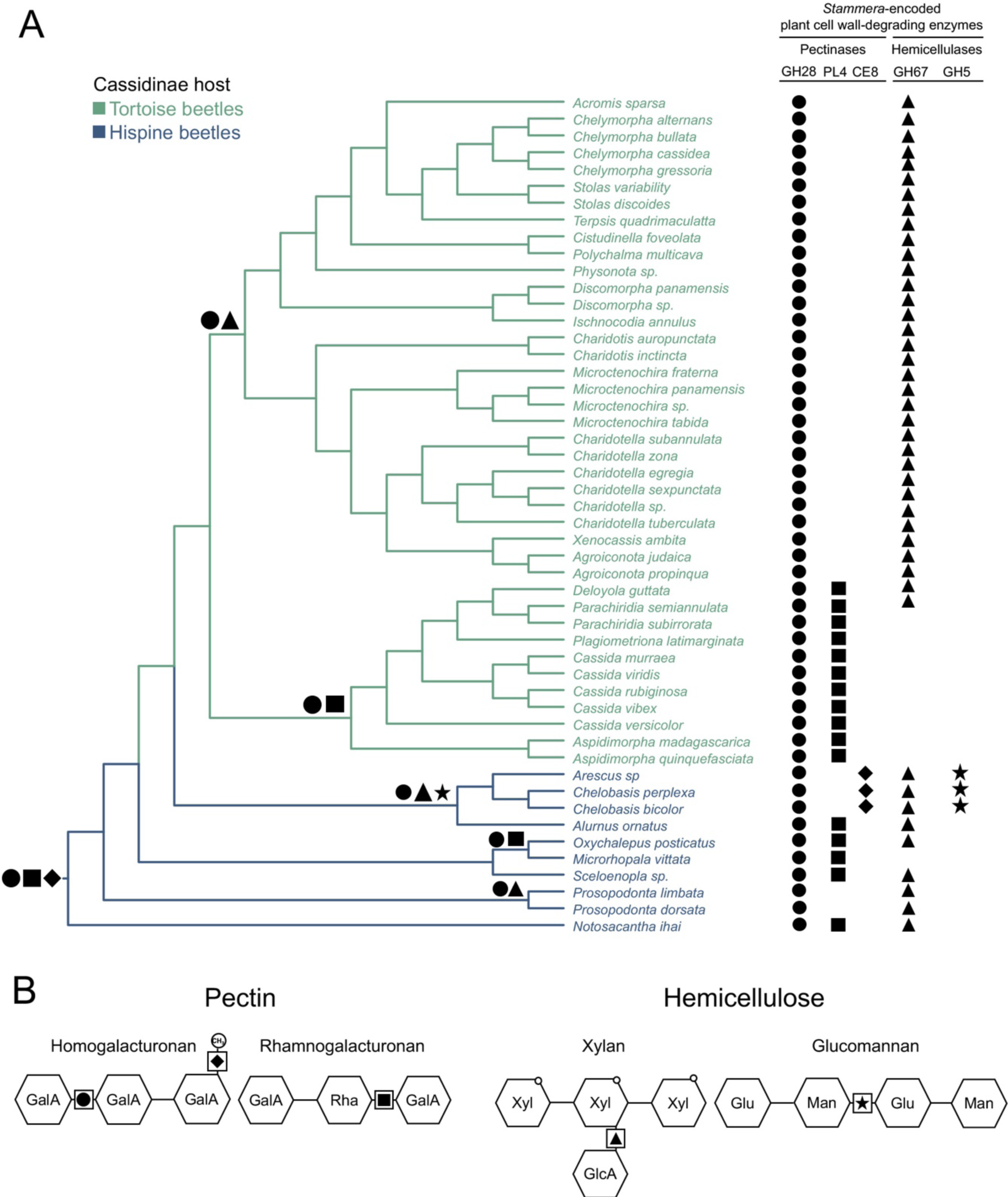
Distribution, assembly, and ancestral reconstruction of *Stammera*-encoded digestive enzymes. (**A**) Distribution of *Stammera* genes encoding plant cell wall-degrading enzymes is represented by different symbols. Circle: GH28 (polygalacturonase), rectangle: PL4 (rhamnogalacturonan lyase) diamond: CE8 (pectin methylesterase), triangle: GH67 (α-glucuronidase) and star: GH5 (endomannanase). Symbiont Maximum Likelihood phylogeny is based on a concatenated alignment of 124 single-copy core genes. The ancestral state of host-beneficial factors was inferred using the trace character history function as implemented in Mesquite v2.75. A character matrix was created for all genes, and likelihood calculations were performed using the Mk1 model. A likelihood score >50% was used to infer ancestral nodes for the different plant cell wall-degrading enzymes encoded by *Stammera* and are illustrated by symbols at the base of each node. Posterior probabilities are summarized in Table S3. (**B**) Predicted mode of action of *Stammera* digestive enzymes across pectin and hemicellulose. Abbreviations: GH, glycoside hydrolase; GlcA, glucuronic acid; Glu, glucose; CE, carbohydrate esterase; Man, mannose; PL, polysaccharide lyase; Rha, rhamnose; Xyl, xylose.

### Ancestral configuration of host-beneficial factors

As the *Stammera*-Cassidinae symbiosis is predicated on the microbe’s ability to deconstruct complex plant polymers ^3,4^, we assessed the distribution, assembly, and ancestral configuration of the symbiont’s plant cell wall-degrading enzymes. Our annotation of *Stammera* genomes spanning tortoise and hispine beetles yielded a diversity of digestive enzymes, matching the divergent nutritional ecology of cassidines (Figure 4). These feature previously-described pectinases such as polygalacturonase (glycoside hydrolase [GH] 28) and rhamnogalacturonan lyase (polysaccharide lyase [PL] 4), together with α-glucuronidase (GH67), a xylanase ^3,4,7^. However, our examination of the partnership beyond tortoise beetles revealed additional *Stammera*-encoded digestive enzymes, including a pectin methylesterase (carbohydrate esterase [CE] 8) and an endomannanase (GH5), both of which are restricted to symbionts of hispines (Figure 4).

Using the trace character history function as implemented in Mesquite ^35^, we reconstructed the distribution of symbiont-encoded digestive enzymes relative to the evolutionary history of *Stammera* (Figure 4). This revealed that the ancestral configuration of the symbiosis featured three plant cell wall-degrading enzymes: polygalacturonase, α-glucuronidase, and rhamnogalacturonan lyase (Figure 4A; Figure S5; Table S3). While pectin methylesterase and endomannanase were acquired secondarily, we observe that gene loss, rather than gain or reacquisition, generally govern the presence of host-beneficial factors (Figure 4A). Such findings are consistent with the metabolic streamlining observed in *Stammera* and other insect endosymbionts ^18,36,37^, where in the absence of opportunities for recombination, gene loss is unidirectional. Except for polygalacturonase, which is encoded by all *Stammera* strains surveyed to date (Figure 4A), four irreversible loss events shaped the distribution of the remaining digestive enzymes (Figure 4A).

Among hispines, the annotation of endomannases in *Stammera* is notable given the predominant specialization of these beetles on monocotyledonous plants ^10^. Monocots represent an especially rich source of glucomannan ^38,39^, and endomannases can deconstruct the hemicellulose by cleaving its β-1,4-linkage of glucose and mannose ^40^ (Figure 4B). As endomannanases are not encoded by *Stammera* in tortoise beetles (Figure 4A), their restriction to symbionts associated with hispines may reflect the ancestral adaptation of cassidines to monocotyledonous plants ^10^, including palms and grasses (Figure 1D). Thus, we predicted that *Stammera* may differentially upgrade the digestive physiology in a subset of hispines against glucomannan relative to tortoise beetles, allowing them to exploit a diet rich in glucomannan. Using thin-layer chromatography, we compared the digestive phenotype of a hispine (*Chelobasis bicolor*) and a tortoise beetle (*Chelymorpha alternans*) bearing metabolically distinct symbionts (Figure S6). Since both species harbor *Stammera* capable of supplementing polygalacturonase, we observe that the two beetles can monomerize homogalacturonan into galacturonic acid (Figure S6A). But confirming *in silico* predictions that *C. bicolor* should depolymerize glucomannan more effectively (owing to its symbiont encoding an endomannanase), we find that the hispine is able to deconstruct the hemicellulose into pronounced dimers and trimers, compared to the breakdown products accumulating in *C. alternans* (Figure S6B).

### Symbiont transcriptional variation matches host nutritional requirements

Characterizing the transcriptional activity of *Stammera* in foregut symbiotic organs of adult beetles revealed a consistent profile that reflects its beneficial role ^7^. Specifically, the gene encoding for polygalacturonase is the 4^th^ most highly expressed transcript, behind ribosomal proteins, but ahead of chaperones such as *groEL*, *groES*, and *dnaK* ^7^. These patterns are in line with the transcriptomes of obligate endosymbionts in other insect groups, where genes coding for chaperones and host-beneficial factors (e.g. essential amino acids) are among the most highly expressed ^41^. But given the symbiont’s extracellular localization within its host and during transmission, ^7^, we aimed to quantify the symbiont’s transcriptional plasticity relative to host development and nutritional requirements. To address this, we compared the transcriptional profiles of *Stammera* within the foregut symbiotic organs of larvae and adults, as well as egg caplets in the tortoise beetle, *C. alternans* (Figure 5A).

**Figure 5.**
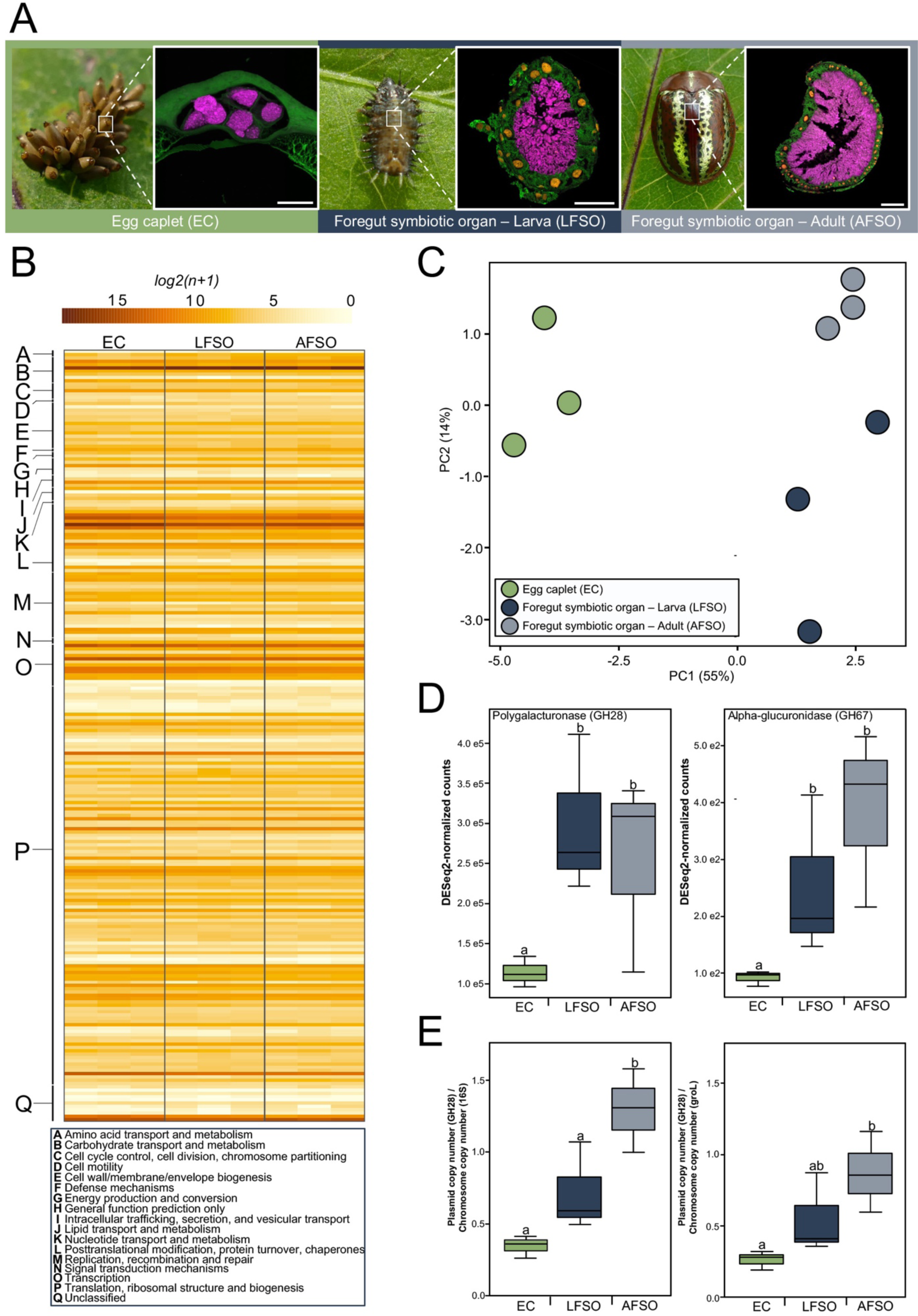
*Stammera* transciptional dynamics match host nutritional requirements. (**A**) *Chelymorpha alternans* eggs, larva (3^rd^ instar), and adult. Fluorescence *in situ* hybridization using cross-sections of egg caplets and foregut symbiotic organs of a larva and an adult targeting *Stammera* (magenta: 16 rRNA) and host (green: 18 rRNA) cells against the DAPI counterstain (yellow). Scale bars (50 μm) are included for reference. (**B**) Heatmap illustrating *Stammera* gene expression across egg caplets and foregut-symbiotic organs in larvae and adults. (**C**) Principal Coordinate Analysis (PCA) of the global transcriptome profile of *Stammera* across host compartments. Significant clustering was assessed by PERMANOVA based on Euclidean distances between samples (*p*=0.003). (**D**) Expression of polygalacturonase (left) and α-glucuronidase (right) genes of *Stammera* across host compartments. Counts were normalized by DESeq2’s median of ratios. Differences in gene expression were calculated using negative binomial generalized linear model (NB-GLM). Different letters above box plots indicate significant differences (polygalacturonase: *X*^2^=14.7, df=2, *p* <0.001; α-glucuronidase: *X*^2^= 25, df=2, *p*<0.005). (**E**) Symbiont plasmid copy number per chromosome across host compartments. This was determined by dividing the polygalacturonase copy number by the copy number of the chromosomal genes: *16S rRNA* (left) and the chaperonin *groEL* (right). Differences in plasmid copy number were estimated using a general linear model (LM) (*16S rRNA*: F_(2,4)_=26.3, df=2, *p*=0.0049, *groEL*: F_(2,4)_=10.37, df=2, *p=*0.0261). Different letters above the box plots indicate significant differences (*p* < 0.05).

Like other cassidines, *C. alternans* relies on egg caplets to vertically transmit *Stammera* ^3,6,8^*. Stammera* is embedded within spherical secretions during transmission and remains separated from the developing embryo until larval eclosion ^8^ (Figure 5A). This contrasts with symbiont localisation within foregut symbiotic organs in larvae and adults (Figure 5A), where *Stammera* is already acquired and is contributing to *C. alternans* digestion and development ^4,8^. Reflecting these differences, we quantified a dynamic transcriptional profile across treatments (Figure 5B, C; PERMANOVA, *p* = 0.003. By comparing the transcriptional activity of *Stammera* within egg caplets relative to larval and adult foregut symbiotic organs (Figure 5B, C), we observe 59 and 65 genes to be differentially expressed, respectively (Dataset 3; FDR-adjusted *p* < 0.05). Most of these genes are shared (52 in total), highlighting a functional overlap in the transcriptional differences affecting *Stammera* during transmission relative to its localization within foregut symbiotic organs (Figure 5C; Dataset 3). In contrast, only 19 *Stammera* genes are differentially expressed between larvae and adult beetles (Dataset 3: FDR-adjusted *p* < 0.05), indicative of a consistent transcriptional profile within symbiotic organs which are morphologically conserved throughout development (Figures 5A-C) ^9^.

Genes that are preferentially expressed in egg caplets relative to the foregut symbiotic organs largely encode chaperones and tolerance proteins (e.g. *cspE*) (Dataset 3; FDR-adjusted *p* < 0.05). These dynamics may reflect the abiotic challenges *Stammera* contends with during extracellular transfer ^17^. On average, *Stammera* must subsist for 11 days within the egg caplet prior to acquisition by eclosing larvae ^8^. Hence, the symbiont may be modulating its metabolism to buffer fluctuations in temperature and humidity ^8,42^, as experienced by other extracellularly-transmitted insect symbionts such as *Ishikawaella* ^43^, *Burkholderia* ^44,45^ and *Pantoea* ^46^.

In contrast, symbiont genes involved in carbohydrate transport and metabolism are upregulated within the foregut symbiotic organs of both larvae and adults relative to the egg caplet (Dataset 3; FDR-adjusted *p* < 0.05), including the two digestive enzymes supplemented by *Stammera* to *C. alternans* (Figure 5D): polygalacturonase (Negative binomial GLM *X*^2^ = 14.7, df=2, *p* < 0.001, average log_2_FoldChange=1.27) and α-glucuronidase (Negative binomial GLM *X*^2^ = 25, df=2, *p* < 0.001, average log_2_FoldChange=1.77). The elevated expression of both genes aligns with the nutritional requirements of larvae and adults since both stages are actively engaged in folivory ^42,47^, in contrast to eggs.

Most endosymbionts possessing drastically reduced genomes (< 0.25 Mb) maintain a rudimentary transcriptional machinery ^34^, limiting their potential to regulate gene expression ^41^. Of the transcription-related genes consistently retained within such tiny genomes, including *Stammera*’s, only *rpoA*, *rpoB*, *rpoC* and *rpoD* are conserved ^34^. But in contrast to intracellular symbionts such as *Carsonella* (0.16 Mb) in psyllids ^48^, *Hodgkinia* (0.14 Mb) in cicadas ^29,49^, and *Nasuia* (0.11 Mb) in leafhoppers ^36^, the extracellular *Stammera* additionally retained a broader collection of transcriptional regulators, featuring *rpoH, rpoZ*, *nusA*, *nusB*, *nusG*, and *rho* (Dataset 4). As major regulators of bacterial transcription elongation, the annotation of *nusA* and *nusG* is especially notable, since both factors modulate intrinsic and Rho-dependent termination by binding to RNA polymerase ^50,51^. A greater transcriptional control may reflect the extracellular localization of *Stammera* (Figure 5A), and the relative instability of life beyond the metabolic comforts of a host cell.

Since polygalacturonase and α-glucuronidase are both plasmid-encoded, we also examined whether *Stammera* can elevate their expression by increasing plasmid abundance. We observe that plasmid copy numbers do increase following larval eclosion from the egg, and through adulthood (Figure 5E; LM, *16S rRNA* [F_(2,4)_=26.3, *p*=0.0049] and *groEL* [F_(2,4)_=10.3, *p*=0.026]. Modulating plasmid abundance to meet host nutritional requirements is shown in other insect-microbe symbioses, including *Buchnera* in aphids ^52^. *Buchnera* genes responsible for the biosynthesis of leucine, an essential amino acid, are encoded on the pLeu plasmid ^53^. In response to leucine starvation in the host, *Buchnera* increases pLeu plasmid copy numbers, mirroring the dynamics observed in *Stammera* relative to beetle development and nutritional requirements (Figure 5E).

### Early-diverging, non-symbiotic cassidines encode polygalacturonase endogenously

As the sole plant cell wall-degrading enzyme universally encoded by *Stammera*, polygalacturonase is critical for origin and stability of symbiosis within the Cassidinae (Figure 4). The glycoside hydrolase similarly underpins the partnership between *Macropleicola* and reed beetles (Chrysomelidae: Donaciinae) ^54^, highlighting the functional convergence of digestive symbioses in folivorous insects ^55^. Given the foundational role of polygalacturonase for leaf beetle-bacterial symbioses ^56^, and the obligate dependence of cassidines on *Stammera* ^3,8^, we aimed to clarify how non-symbiotic members of the subfamily contend with a leafy diet enriched in pectin.

In addition to symbiosis, horizontal gene transfer from bacteria and fungi endowed herbivorous beetles with catalytic tools to deconstruct complex plant polymers ^2,57^. The two independent origins of herbivory in beetles coincided with the cooption of microbial plant cell wall-degrading enzymes ^2^, including polygalacturonase ^57^. And while symbiotic cassidines maintain cellulases endogenously, beetle-encoded polygalacturonases were lost ^4^, suggesting that the acquisition of *Stammera* may have relaxed selection to retain the enzyme. To explore whether early-diverging, non-symbiotic members of the subfamily encode the pectinase endogenously, we characterized the plant cell wall-degrading enzymes maintained by *Calyptocephala attenuata*, a hispine beetle belonging to the Spilophorini tribe (Figure 1B). By combining long-read High Fidelity (HiFi) genome sequencing from Pacific Biosciences with RNA sequencing on an Illumina NextSeq 2000 system, we observe that *C. attenuata* encodes the same set of cellulases (GH9, 45 and 48) and xylanases (GH10) as symbiotic members of the Cassidinae (Figure 6). But in contrast, we also annotated polygalacturonase-encoding genes on three separate beetle contigs (Figure 6A), ranging in size between 25 and 45 Kb and flanked by insect genes (Figure S7). Unlike ancestral polygalacturonases across the Chrysomelidae, beetle encoded GH28 by *C. attenuata* most closely align with pectinases from Betaproteobacteria (Figure S10). Intron content and TATA box localization further clarified the eukaryotic features of these genes, coupled with our ability to amplify them by PCR using DNA extractions from the legs, thorax, and elytra of *C. attenuata* (Figure S8). All three copies of the gene retained the key catalytic residues described for polygalacturonase (Figure 6A) (Figure S9) ^58,59^, indicating that the encoded enzymes are functionally conserved and likely confer a pectinolytic phenotype in the absence of *Stammera*.

**Figure 6.**
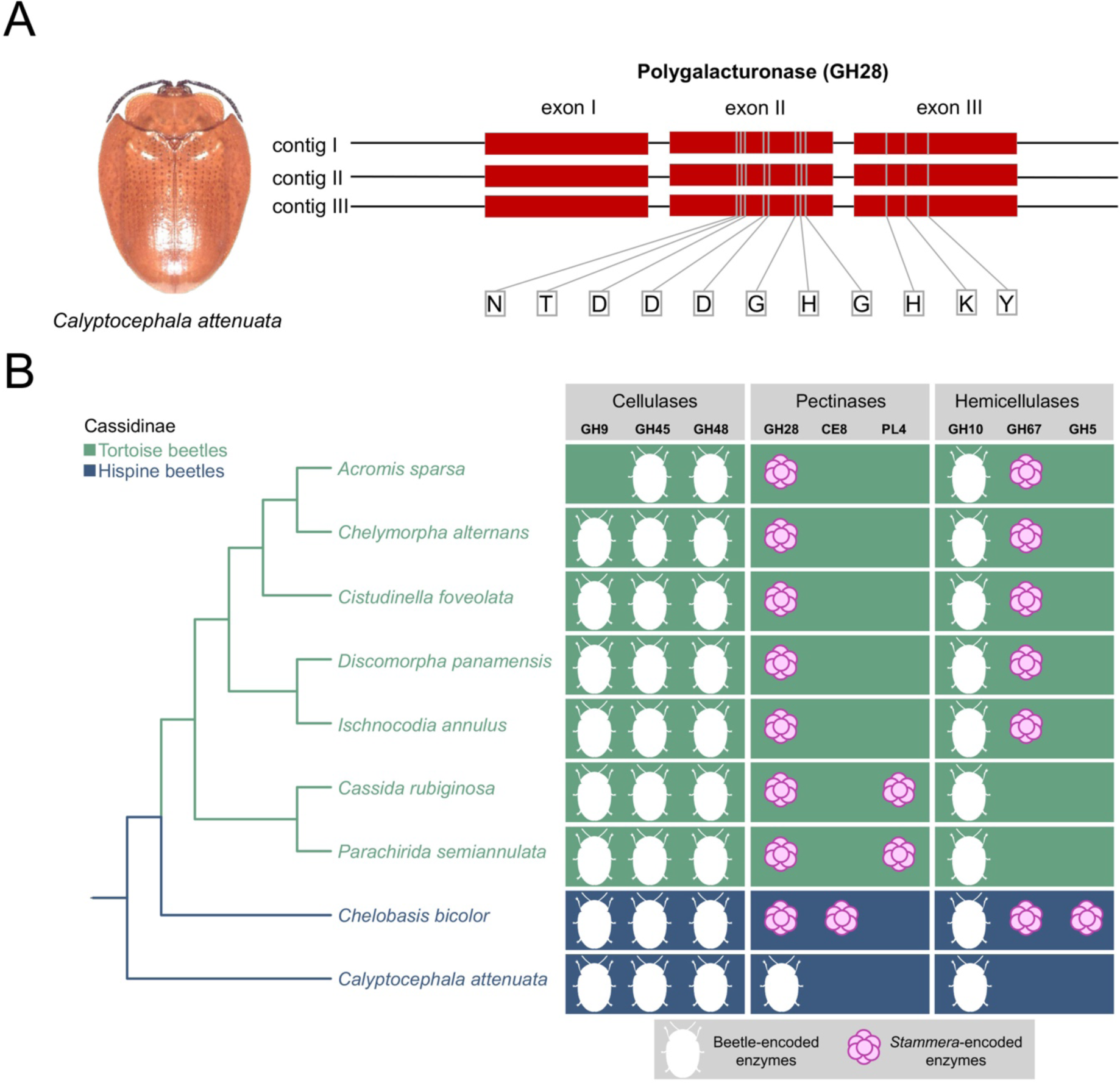
Early-diverging, non-symbiotic cassidines encode polygalacturonase endogenously. (**A**) The early-diverging, non-symbiotic hispine beetle *Calyptocephala attenuata.* Gene structure and functionally conserved amino acids across the three copies of the polygalacturonase-encoding gene from *C. attenuata* draft genome assembly. (**B**) Distribution of host- and *Stammera*-encoded plant cell wall-degrading enzymes as inferred from transcriptome and symbiont genome profiling of nine representative cassidine species. Source of each digestive enzyme is designated by an icon. Abbreviations: GH, glycoside hydrolase; CE, carbohydrate esterase, PL, polysaccharide lyase.

Given the adaptive importance of polygalacturonase for herbivorous beetles ^57,60^, it is unclear how selection may have favored its outsourcing to *Stammera* in symbiotic cassidines (Figure 6B) ^3,4^. It is conceivable that deleterious mutations may have compromised the functionality of beetle-encoded pectinases, necessitating rescue through symbiosis with a beneficial microbe. Several herbivorous beetles retain a repertoire of functionally active and inactive polygalacturonases ^57,61^, where the latter can no longer bind homogalacturonan due to amino acid substitutions at crucial positions ^62^. It is also possible that symbiont acquisition may have spurred evolutionary innovation by upgrading the digestive abilities in a subset of cassidines, rendering endogenous polygalacturonases redundant. Both scenarios are not mutually exclusive and may have occurred in a stepwise process. While non-symbiotic cassidines retained polygalacturonase endogenously (Figure 6B), the ancestral configuration of enzymes derived from *Stammera* included polygalacturonase in addition to pectin methylesterase, endomannanase and α-glucuronidase (Figure 4A). Convergently, the acquisition of the pectinolytic symbiont *Macropleicola* by reed beetles also coincided with the loss of host-encoded polygalacturonases ^54^. While speculative, by expanding the range of universal plant polymers that an herbivore can deconstruct (Figure 4), *Stammera* may have relaxed selection for its host to maintain polygalacturonase and/or compensated for their reduced efficiency.

### Beetle diversification following symbiont acquisition

Symbioses are key drivers of global biodiversity ^63,64^. By facilitating access to new environments, or by allowing organisms to integrate novel metabolic features, mutualistic partnerships can promote diversification by increasing speciation rates, and/or by decreasing the rate of extinction ^63,64^. The consequences of beneficial partnerships on species richness are most evident when net diversification rates are compared between symbiotic and nonsymbiotic members of a clade. In gall-inducing midges, for example, the acquisition of a fungal nutritional symbiont resulted in a sevenfold expansion in the range of suitable host-plant taxa relative to lineages that do not stably associate with a fungus ^65^. Correspondingly, net diversification of symbiotic midges outpaced that of their non-symbiotic relatives by 17 times ^65^.

Among cassidines, the loss of endogenous polygalacturonases coincided with the acquisition of a symbiont supplementing a broader collection of pectinases and other digestive enzymes (Figure 6B). Given the expanded metabolic potential of symbiotic cassidines relative to non-symbiotic taxa, we quantified how *Stammera* acquisition impacted the diversification rate, species richness, and plant use by its host. Three fossil calibration points were used to infer the minimum ages of divergence within the Cassidinae phylogeny, including Notosacanthini, Chalepini, and Cassidini specimens dated at 47 ^66^, 44.1 ^11^, and 40 Mya ^67^, respectively. The resulting time-calibrated phylogeny revealed that the symbiosis was formed 62 Mya (95% CI: 59.9 to 64.3), soon after the Paleocene origin of the Cassidinae subfamily (62.5 Mya; 95% CI: 59.36 to 65.59) (Figure 7; Figure S11; Table S4). By applying estimates of tribe-level species richness within the Cassidinae ^10^, net diversification rates were quantified relative to the symbiotic condition using MEDUSA and as implemented in the Geiger package ^68^. Our analysis revealed a background speciation rate of 0.0557 (lineages/Myr) and located two diversification shifts (Figure S12). A net increase rate followed *Stammera* acquisition (Figure S12; 0.1237 lineages/Myr), potentially implicating symbiosis in the ecological radiation of the Cassidinae subfamily and its accumulation of species diversity (Figure 7). This is concordant with our comparison of species richness and host-plant use of symbiotic and non-symbiotic cassidines. We observe that *Stammera*-harboring tribes are significantly more speciose (Figure 7) (G-test, G = 182.14, df=1, *p* < 0.001) and exploit a greater variety of plant families (G-test, G = 43.64, df = 1, *p* < 0.001). A second, and more derived, diversification rate shift featured a slowdown in two symbiotic hispine clades, the Alurnini and Arescini (Figure S11; 0.0594 lineages/Myr), Arescini feature specialists on the immature rolled leaves of plants in the monocotyledonous genus *Heliconia* ^12^. McKenna and Farrell ^11^ report similar decelerations in *Heliconia*-feeding hispine beetles, suggesting that niche specialization may decrease origination rates among cassidines irrespective of an association with *Stammera*.

**Figure 7.**
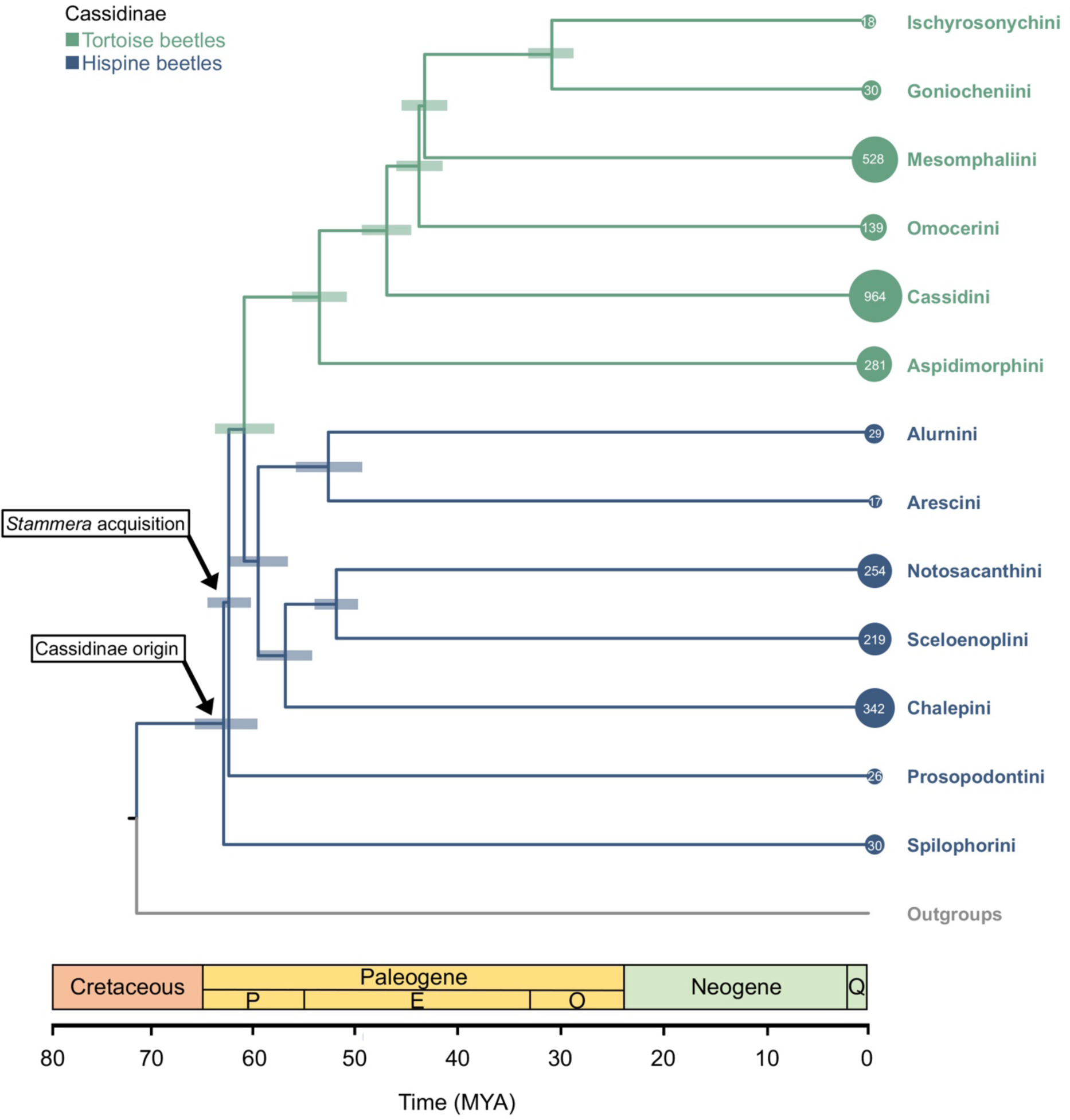
Symbiont acquisition relative to the evolutionary history of Cassidinae beetles. Time-calibrated phylogeny dating the origin of the Cassidinae subfamily and the timing of *Stammera* acquisition. Branches are coloured to differentiate hispine (dark blue) from tortoise beetle tribes (light blue). Circle sizes (and their enclosed numbers) correspond to the species richness of each Cassidinae tribe. Bars depict confidence intervals (95% highest posterior density) of node ages. Abbreviations: P, Paleocene; E, Eocene; O, Oligocene; Q, Quaternary.

## Conclusions

Symbioses evolved across several highly diverse insect clades in conjunction with key traits to facilitate herbivory ^37,56,69–72^. Through the supplementation of nutrients to balance a specialized diet ^36,72,73^, or the production of enzymes to overcome complex plant molecules and toxins ^70,74–76^, symbiont acquisition and evolution are key determinants of host-plant use ^77^. Despite marked differences in their nutritional ecology, here we report on a shared symbiosis between tortoise and hispine beetles, offering insights into the origin and ancestral configuration of a Paleocene-age partnership with *Stammera*. In light of an upgraded metabolic potential following symbiont acquisition, *Stammera*-harboring cassidines are more speciose and exploit a greater variety of plants, highlighting the adaptive impact of a symbiotic transition.

## Methodology

### Taxon sampling

Adult Cassidinae species were collected in France, Germany, Japan, New Zealand, Panama, and the United States of America between 2018-2023. For DNA sequencing, the insects were submerged in molecular grade 99% ethanol after collection and kept at -20 °C for further processing. *Calyptocephala attenuata* and *Chelobasis bicolor* were snap frozen in liquid nitrogen ahead of RNA extraction and transcriptome sequencing. *C. bicolor* was also used for enzymatic assays, along with *Chelymorpha alternans*, which is continuously reared at the Max Planck Institute for Biology (Tübingen, Germany). The latter species was also used to study *Stammera*’s gene expression across different host developmental stages and compartments. All experiments were performed in accordance with relevant guidelines and are in compliance with EU and German legislation on insect rearing and experimentation.

### Genome sequencing and assembly

Metagenomic sequencing was performed across 55 representative Cassidinae beetle species and spanning 13 tribes. 42 of these species were collected in this study and dissections were preformed using 1-3 individuals under molecular grade 99% ethanol. DNA was extracted using the QIAGEN DNeasy Blood & Tissue Kit with RNase treatment. Genomic DNA was fragmented to an average size of 300 bp using Covaris S2. Sheared DNA was purified by SPRI beads and used to construct DNAseq libraries using the NEBNext® Ultra™ II DNA Library Prep Kit for Illumina. 12 libraries were sequenced on an Illumina HiSeq 3000 system, and due to an update of Illumina sequencing technologies at the institute, 30 libraries were sequenced on an Illumina NextSeq 2000 system at the Max Planck for Biology (Tübingen, Germany) using the paired-end 150 bp technology with a depth of ∼50 million reads.

Adaptor removal and quality filtering of raw reads was performed by Trimmomatic (v0.36) ^78^. Metagenomic sequences of 42 Cassidinae species generated in this study in addition to 13 publicly available Cassidinae sequencing read sets ^4^ were *de novo* assembled by MEGAHIT ^79^. Contigs belonging to *Stammera* were binned according to coverage and GC content by CONCOCT ^80^. Out of the 50 *Stammera* genomes, 27 were automatically assembled into a single chromosomal sequence, while the other 23 consisted of 2 to 8 contigs. These contigs were reordered and scaffolded by comparing them to a complete *Stammera* assembly from the same host genus using the Mauve Aligner ^81^. Subsequently, genome curation procedures, involving the filling of scaffolding gaps in the 23 fragmented genomes and the removal of local assembly errors in all 50 genomes, were carried out following established protocols^82^. After verification of complete single-chromosomal *Stammera* genomes, GC skew was calculated by iRep ^83^ to identify the origin of replication. OriC sites were then set in Geneious Prime 2019.2.3 (https://www.geneious.com).

### *Stammera* comparative genomics Symbiont genome annotation

Symbiont protein-coding genes were predicted by Prodigal as implemented in Prokka (v1.14.6)^84^ using the genetic code 4 (TGA encoding tryptophan) as described by Salem *et al* ^4^. Additional gene predictions were performed by Glimmer^85^ and a manual curation of annotation files was performed to consolidate the gene predictions. tRNAs, rRNAs and ncRNAs were predicted by ARAGORN, Barrnap and Infernal as implemented in Prokka (v1.14.6) ^84^. Pseudogenes were predicted in each genome using Pseudofinder^86^.

### Pangenome analysis

Pfam, KOfam, NCBI COG’s, and KEGG annotations were additionally included in anvi’o (v8.1-dev)^24^. Subsequently, a pangenome analysis of 50 *Stammera* genomes was performed (minbit= 0.3, MCL inflation parameter=2) using DIAMOND^87^ for amino acid sequence similarity search (--sensitive) ^24^. Orthologous genes were considered part of the core genome if present in 100% of the genomes. Correlation between *Stammera*’s genome size and number of protein-coding genes was evaluated using a Spearman’s rank correlation test in R v. 4.3.1 ^88^. To compare *Stammera* genomes at the nucleotide level, the Average Nucleotide Identity (ANI) between symbiont genomes was calculated using pyani ^89^ as implemented in anvi’o. A Pearson’s X^2^ test was used in R v. 4.3.1 ^88^ to determine *Stammera* genome relatedness within and between tortoise and hispine beetles. Conservation of gene order in Stammera genomes was assessed by MCScanX, including a representative of each tribe ^90^, and the results were visualized in SynVisio ^91^. Distribution of gene clusters across *Stammera* from tortoise and hispine beetles was visualized by an Upset plot based on a presence/absence matrix obtained from *<u>Stammera</u>*’s pangenome. This plot was constructed using the R packages UpSet ^92^ and ComplexHeatmap ^93^. A pangenome with functions was obtained with the program anvi-display-functions in anvi’o to compare gene functions rather than gene sequences across *Stammera* genomes, resulting in a frequency and a presence/absence table of COG categories.

### Symbiont phylogenetic reconstruction

124 single-copy core genes identified by anvi’o were extracted from each *Stammera* genome and individually aligned using MUSCLE (v3.8.1551) ^94^. Unrooted phylogenies encompassing the 50 *Stammera* taxa were constructed from a concatenated multigene alignment by Bayesian and Maximum Likelihood (ML) methods using MrBayes (v3.2.7a) ^95^ (ngen=1000000, samplefreq=1000) and RAxML-NG (v1.2.0) ^96^ (ngen=1000), respectively (Figure S13). Concatenated alignments of 61 single-copy genes present in all *Stammera* taxa, and 26 outgroups indicated in Table S5, were included to construct Bayesian and ML phylogenies using *Xanthomonas campestris* as a root (Figure S3). The best-fit substitution models for each analysis were selected using PartitionFinder2 ^97^ (branchlengths=unlinked, models = all, model_selection = bic) (Table S6).

### *Stammera* molecular evolution

To determine the signatures of selection acting on *Stammera’*s genes, we measured rates of synonymous (d*S*) and nonsynonymous (d*N*) substitutions across orthologous single-copy genes. This parameter determines whether genes experience strong purifying selection (ω < 0.1), relaxed purifying selection (1 < ω > 0.1), or positive selection (ω > 1) ^26^. Codon-based alignments were performed for each gene in PAL2NAL (v14) ^98^ and by using the unrooted *Stammera* Bayesian tree. d*N*/d*S* ratios were estimated for each using codeml as implemented in PAML (v4.9) ^31^ by applying three models. The M0 model was used to test for selection across all codon sites. Additionally, the site-based model M1a (nearly neutral) allowing for two categories of sites (ω = 1 and ω = 0), was compared to the M2a model (positive selection) which allows an additional category of positively selected sites (ω >1). To test for sites with significant support for each model, Likelihood Ratio Tests (LRTs) were compared against *X*^2^.

### Ancestral state reconstruction

Genes encoding for plant-cell wall digestive enzymes were identified in *Stammera*’s pangenome. Ancestral nodes of these genes were inferred in the unrooted Maximum Likelihood *Stammera* tree using the trace character history function as implemented in Mesquite (v.3.7) ^35^. This phylogeny was rooted according to the *Stammera* tree that included the outgroup species indicated in Table S5. A category character matrix was created using a gene presence/absence table and likelihood calculations were performed using the Mk1 model. Ancestral nodes for each gene were identified in the symbiont tree using a cut-off likelihood value > 50%.

### Host phylogenetic reconstruction

Host mitochondrial genomes were extracted from metagenomic assemblies based on coverage and GC content. BLAST searches confirmed the insect origin of mitochondrial genomes and these were further annotated using the MITOS2 webserver (http://mitos2.bioinf.uni-leipzig.de) ^99^. A concatenated alignment of 15 mitochondrial genes (13 protein coding genes + 2 ribosomal rRNA genes) was partitioned to assign the most appropriate substitution model to each gene using PartitionFinder2 (Table S6). Phylogenetic analyses were performed using both Bayesian inference and Maximum Likelihood in MrBayes ^95^ (ngen=5000000, samplefreq=1000) and RAxML-NG ^96^ (n=1000) , respectively (Figure S13). Members of Spilopyrinae and Eumolpinae subfamilies from the Chrysomelidae were used as outgroups for this analysis. Bayesian time calibrated phylogenies were inferred by BEAST2 (v2.4.8) ^100^ using the generated partition scheme. Substitution models were selected by bModelTest as implemented in BEAST. The tree prior included the calibrated Yule Model with a random starting tree. Three internal node calibrations, Notosacanthini (47 Mya) ^66^, Chalepini (44.1 Mya) ^11^, and, Cassidini (40 Mya) ^67^, were applied with a normal prior distribution. Multiple BEAST chains (ngen=10000000) were run per genome alignment and sampled every 1000 generations with a strict clock mode.

### Host-symbiont cophylogenetic analysis

The tree reconciliation software eMPRess GUI ^101^ was used to study the evolutionary relationship between Cassidinae and *Stammera*. This software reconciles symbiont and host trees using the DuplicationTransfer-Loss (DTL) model. Host and symbiont Maximum Likelihood phylogenetic trees were used as input and the analysis was conducted using the following eMPRess parameters: duplication cost = 1, transfer cost = 1, and loss cost = 1. The significance of reconciliation between host and symbiont tree was calculated by randomizing the tips of the branches and then, re-calculating the cost to reconcile the phylogenies. Congruent phylogenies are obtained when the original cost of reconciliation is less than expected by chance (*p*<0.01).

### Diversification rate analyses

We applied MEDUSA (Modeling Evolutionary Diversification Using Stepwise Akaike Information Criterion) ^102^ to estimate shifts in the diversification of the Cassidinae relative to *Stammera* acquisition. For this analysis, we collapsed the time-calibrated Cassidinae phylogeny obtained from BEAST to incorporate tribe-level species estimates reported by Chaboo ^10^. As a complementary test, we compared species richness between non-symbiotic and symbiotic cassidines using the G-test of goodness-of-fit ^103^. We also investigated whether symbiotic beetles exploit greater diversity of plant families relative to non-symbiotic cassidines. Host-plant family assignments for each Cassidinae tribe was also accessed from Chaboo ^10^. As outlined by Edger et al ^104^, G-tests of goodness-fit were performed in R v. 4.1.1 ^88^ using the DescTools package ^105^ to test if our observed values are significantly different from expectations, assuming equal species in both conditions. A Williams correction was implemented for a better approximation of the chi-square distribution, resulting in a more conservative test ^103^.

### Symbiont transcriptome sequencing

To characterize differences in symbiont gene expression relative to *Stammera* localization and host development, transcriptome sequencing of egg caplets, foregut symbiotic organs of 3^rd^ instar *Chelymorpha alternans* larvae and 24-day-old adults was performed. Six egg clutches of ∼ 30 eggs representing three replicates were first divided in half two days after oviposition. Caplets were removed with sterilized scissors from one of each half. Treatments were pooled, yielding 30 caplets in each replicate. The remaining eggs were maintained under standard growth conditions (26°C and 60% relative humidity) previously described in Pons *et al* ^8^ and Berasategui *et al* ^106^. Three larvae from each replicate were collected nine days after hatching. Foregut symbiotic organs were dissected from larvae using sterilized scissors. The remaining larvae were kept under the same conditions until reaching adulthood. Three female adults from each replicate were sampled 24 days after emergence and foregut symbiotic organs were dissected for further processing. RNA extraction was performed for each sample immediately after collection using the QIAGEN RNeasy Mini Kit according to the protocol 4: Enzymatic Lysis and Proteinase K Digestion of Bacteria starting from step 7. This protocol is included in the RNAprotect® Bacteria Reagent Handbook from Qiagen. Total RNA was further quantified using the Qubit™ RNA High Sensitivity (HS) kit. 30 ng of total RNA were used as input to prepare nine RNA sequencing libraries for egg caplets, foregut symbiotic organs of larvae, and foregut symbiotic organs of adults (3 biological replicates each). Due to sequence similarity, ribosomal RNA was depleted from total RNA using the NEBNext® rRNA Depletion Kit (Human/Mouse/Rat). Libraries were constructed using the NEBNext® Ultra™ Directional II RNA Library Prep and their size was confirmed in a 2100 Bioanalyzer system using the Agilent Technologies High Sensitivity DNA Kit. Sequencing of final libraries was performed on an Illumina HiSeq 3000 system (2×150bp) at the Max Planck Institute for Biology (Tübingen, Germany) with a depth of 30 million reads.

Adapter removal and quality filtering of raw reads was performed in Trimmomatic ^78^. Filtered reads were mapped to *Stammera* genome using bowtie2 (v2.3.5.1) ^107^ (--fr, --no-unal). Gene counts were summarized using the featureCounts program ^108^ (--p, --countReadPairs, --s 2) as part of the Subread package release 2.0.0. Read counts were normalized by the DESeq2’s median of ratios as established in the DESeq2 ^109^ package in R. A heatmap of the log(x+1) normalized reads was then constructed using the pheatmap package ^110^ to visualize the expression profile across samples. Global transcriptome profile between samples was compared by testing for significant clusters using a permuted multivariate analysis of variance (PERMANOVA) using the vegan::adonis() function in R v.4.3.1 ^111^. The likelihood ratio test (LRT), implemented in DESeq2, was used to test for differences in symbiont gene expression across egg caplets and foregut-symbiotic organs of larvae and adults. Differentially expressed symbiont genes were identified with the following criteria: adjusted FDR < 0.05 and fold-change > 0.5. To compare the expression of polygalacturonase and α-glucuronidase genes of *Stammera* across host compartments, normalized transcripts were analyzed using a negative binomial generalized linear model implemented by the ‘glm.nb’ function of the R (4.3.1) package *MASS* ^112^. *Post hoc* Tukey HSD test was performed using the ‘glht’ function of the R package *multcomp* ^113^ with Bonferroni corrections.

### Quantitative PCR

Symbiont plasmid copy number was measured across host compartments by quantitative polymerase chain reaction (qPCR). DNA was extracted from egg caplets, foregut symbiotic organs of larvae and adults using the QIAGEN DNeasy Blood & Tissue Kit with RNase treatment. PCR reactions of 25μl were set up using the Qiagen SYBR Green Mix using the following parameters: 95°C for 10 min, 45 cycles of 95°C for 30 s, 62.7°C for 20 s, and a melting curve analysis was conducted by increasing temperature from 60 to 95°C during 30 s on an Analytik Jena qTOWER³ cycler. Standard curves (10-fold dilution series from 10^−1^ to 10^−8^ ng μl^−1^) were generated using purified PCR products. Absolute gene copy numbers were obtained by interpolating the obtained Ct value against the standard curve. Plasmid copy number was determined by dividing the polygalacturonase copy number, localized in plasmids, by the absolute copy number of the chromosomal genes chaperonin GroEL (*groEL*) and the *16S rRNA* gene (Primers in Table S7). Differences in plasmid copy number across host compartments were analyzed using a general linear model, after validation of a normal distribution, and using host compartments and replicates as fixed factors. *Post hoc* Tukey HSD test was performed using the ‘glht’ function of the R package *multcomp* with Bonferroni corrections in R v.4.1.1 ^111^.

### Long-read sequencing and beetle genome assembly

High molecular weight genomic DNA was extracted from the whole body of four *Calyptocephala attenuata* adults using the Monarch HMW DNA extraction kit for tissue from NEB. The extracted DNA was sheared to between 15 kb and 20 kb using the Megaruptor 2 (Diagenode). A HiFi sequencing library was prepared using SMRTbell Express Template Prep Kit 2.0. Size selection of the final library was performed by the BluePippin System from SAGE Science. Fractions for sequencing were selected based on results from the Femto Pulse System. Desired size fractions were pooled and the final library was purified and concentrated using AMPure PB beads. Quantity of the final library was assessed using the Qubit™ dsDNA HS Assay Kit and the final size distribution was confirmed on the Femto Pulse. Sequencing was performed using one 8M SMRT cell on the PacBio Sequel II System at the Max Planck for Biology (Tübingen, Germany). The pbbccs tool from the pbbioconda package (–min-passes 3 –min-rq 0.99 –min-length 10 –max-length 50000) was utilized to generate the High Fidelity (HiFi) reads (>Q20). Genome assembly was performed by Hifiasm (v0.14.1-r314) ^114^. Polygalacturonase gene identified initially with Illumina sequencing was aligned to the *C. attenuata* draft assembly. Polygalacturonase-containing contigs were extracted and further annotated by AUGUSTUS ^115^ using *Tribolium* as a training set. PCR targeting this gene in legs, thorax and elytra of *C. attenuata* confirmed that early-diverging cassidinaes from the Sphilophorini tribe encode a polygalacturonase gene endogenously (PCR primers Table S8).

### Host RNAseq, transcriptome assembly, and CAZy annotation

Internal organs from *Chelobasis bicolor* and *Calyptocephala attenuata* were dissected and snap-frozen in liquid nitrogen. Total RNA was extracted using the QIAGEN RNeasy Mini Kit with DNase treatment. RNA was quantified using the Qubit™ RNA High Sensitivity (HS) kit. 600 ng of total RNA were used as input for RNA sequencing library preparation. mRNA enrichment was performed using the NEBNext® Poly(A) mRNA Magnetic Isolation Module. RNAseq libraries were constructed using the NEBNext® Ultra™ Directional II RNA Library and their size was confirmed in a 2100 Bioanalyzer system using the Agilent Technologies High Sensitivity DNA Kit. Sequencing was performed on an Illumina NextSeq 2000 system at the Max Planck for Biology (Tübingen, Germany) using paired-end chemistry (2×150bp) with a depth of ∼40 million reads. Adapters were removed from reads and quality filtered by Trimmomatic ^78^. RNAseq reads for *Acromis sparsa*, *Chelymorpha alternans*, *Cassida rubiginosa*, *Parachiridia semiannulta*, *Ischnocodia annulus*, *Cistudinella foveolata,* and *Discomorpha panamensis* were retrieved from NCBI (accession numbers in Table S9) and included in this comparative analysis. *De novo* transcriptome assemblies for nine cassidines was performed using the Trinity platform (v2.8.5) ^116^ (--normalize_by_read_set, --SS_lib_type RF). Assemblies were assessed by BUSCO (v5.1.2) ^117^ using the OrthoDB v.10 Endopterygota gene set ^118^ (Table S9). Protein-coding genes were identified from transcriptome assemblies by TransDecoder (v5.5.0) ^119^. Carbohydrate-active enzymes (CAZys) present were annotated by the dbCAN2 standalone tool run_dbcan4 ^120^.

### Thin layer chromatography (TLC)

Qualitative analysis of breakdown products was performed by thin layer chromatography (TLC) of 20 μl enzyme assays set up as follows: 14 μl of crude gut extract of symbiotic *Chelymorpha alternans* and *Chelobasis perplexa* were incubated with 0.2% polygalacturonic acid in 20 mM citrate/phosphate buffer pH 5.0 at 40°C for 16 h. Polygalacturonase from *Aspergillus niger* was used as a positive control (16 μl of a 0.08 mg/ml solution). Incubated samples were further diluted (1:4) with H_2_O and a total of 16 μl were applied to TLC plates (Silica gel 60, 20 × 20 cm, Merck) in 4 μl steps. Plates were ascendingly developed with ethyl acetate:glacial acetic acid:formic acid:water (9:3:1:4) for about 90 min. After drying, carbohydrates were stained by dipping the plates in a solution containing 0.2% (w/v) orcinol in methanol:sulfuric acid (9:1), followed by a short heating until spots appeared. The reference standard contained 1μg/μl each of galacturonic, di-galacturonic and tri-galacturonic.

Endomannanase activity was qualitatively assessed using 100 μl enzyme assays set up as follows: 40 μl crude gut extract of *Chelymorpha alternans* and *Chelobasis perplexa* were incubated with 0.15% Glucomanan in 30 mM citrate/phosphate buffer pH 6.0 at 70°C for about 1h. Mannanase from *Aspergillus niger* was used as a positive control (1 μl of a 10μg/ml solution). A total of 16 μl from each sample were applied to TLC plates (Silica gel 60, 20 × 20 cm, Merck) in 4 μl steps. Plates were developed ascending with 1-Butanol:glacial acetic acid:water (2:1:1) for about 90 min. After drying, carbohydrates were stained by dipping the plates in a solution containing 0.2% (w/v) orcinol in methanol:sulfuric acid (9:1), followed by a short heating until spots appeared. The reference standard contained 1μg/μl each of mannose, mannobiose, mannotriose and mannotetraose.

### Fluorescence *in situ* hybridization (FISH)

To localize *Stammera* in foregut-symbiotic organs of hispine and tortoise beetle species, as well as in eggs and foregut-symbiotic organs of *C. alternans* larvae and adults, we applied fluorescence *in situ* hybridization (FISH) on paraffin sections. Eggs and foregut-symbiotic organs were dissected and fixed in 4% paraformaldehyde (paraformaldehyde: PBS1X) (v/v) at room temperature during 4h under gentle shaking (500 rpm). After dehydration in an increasing ethanol series of 50, 70, 80, 96 and 100% (v/v) for 1h each, samples were further dehydrated in Roti®-Histol (Carl-Roth, Germany) overnight and embedded in paraffin (Paraplast High Melt, Leica, Germany) overnight. The paraffin-embedded samples were cross-sectioned at 10 µm using a microtome and mounted on poly-L-lysine-coated glass slides (Epredia, Germany) in a water bath. Paraffin sections were left to dry in vertical position at room temperature overnight and baked at 60°C for tissue adherence improvement. They were dewaxed with Roti®-Histol in three consecutive steps for 10 min each followed by decreasing ethanol series of 100, 96, 80, 70 and 50% (v/v) for 10 min each and then washed in milliQ water for 10 min. Slides were dried at 37°C for 30 min and sections were surrounded by a PAP-pen circle to avoid buffer leaking during hybridization. Foregut-symbiotic organs of Cassidinae beetles were hybridized with the probe SCA600 doubly labeled with the fluorophore Cy5 whereas eggs and foregut-symbiotic organs of *C. alternans* larvae and adults were hybridized with the probe SAL227 doubly labeled with the fluorophore Atto550. The oligonucleotide probe SCA600 was designed to target the 16S rRNA sequence of *Stammera* from 52 cassidines and the SAL227 probe was designed to specifically target the 16S rRNA sequence of Stammera from *C. alternans* using the using the software ARB (99) (Table S10). To target host tissues, the generic eukaryotic probe EUK-1195 (Table S10) doubly labeled with the fluorophore Atto488 was included in both hybridization treatments. All probes were dissolved at 5 ng µl-1 in the hybridization buffer containing 35% formamide (v/v), 900 mM NaCl, 20 mM Tris-HCl pH 7.8, 1% blocking reagent for nucleic acids (v/v) (Roche, Switzerland), 0.02 SDS (v/v) and 10% dextran sulfate (w/v). Fifty microliters of hybridization buffer were used per section. The slides were placed in a hybridization chamber at 46°C for 4h with KIMTECHScience precision wipes (Kimberly-Clark, TX, USA) partially soaked in formamide 35% to maintain a humid atmosphere. Sections were rinsed in pre-warmed 48°C washing buffer (70 mM NaCl, 20 mM Tris-HCl pH 7.8, 5 mM EDTA pH 8.0, and 0.01% SDS (v/v)) and transferred to fresh pre-warmed washing buffer for 15 min followed by 20 min in room temperature 1X PBS and 1 min in room temperature milliQ water. After washing, sections were counterstained with DAPI for 10 min at room temperature, dipped in milliQ water, dipped in ethanol 100% and dried at 37°C for 30 min. Slides were mounted using the ProLong® Gold antifade mounting media (Thermo Fisher Scientific, MA, USA), cured overnight at room temperature and stored at -20°C until visualization. Samples were visualized using an LSM 780 confocal microscope (Zeiss, Deutschland). All steps during and after hybridization were done in darkness.

## Acknowledgments

We thank Julie Johnson (Life Sciences Studios) for the illustrations in Figure 1D. Financial support from the Max Planck Society, German Research Foundation (SA 3105/2-1, BE 6922/1-1, and EXC2124—390838134), Japan Science and Technology Agency (JST) (ERATO grant JPMJER1902), and Alexander Von Humboldt Foundation is gratefully acknowledged.

