## Supplementary information for "Paleocene origin of a streamlined digestive symbiosis in leaf beetles"

**This PDF file includes:**

Supporting text

Figures S1 to S12

Tables S1 to S10

SI References

**Other supporting materials for this manuscript include the following:**

Datasets S1 to S4

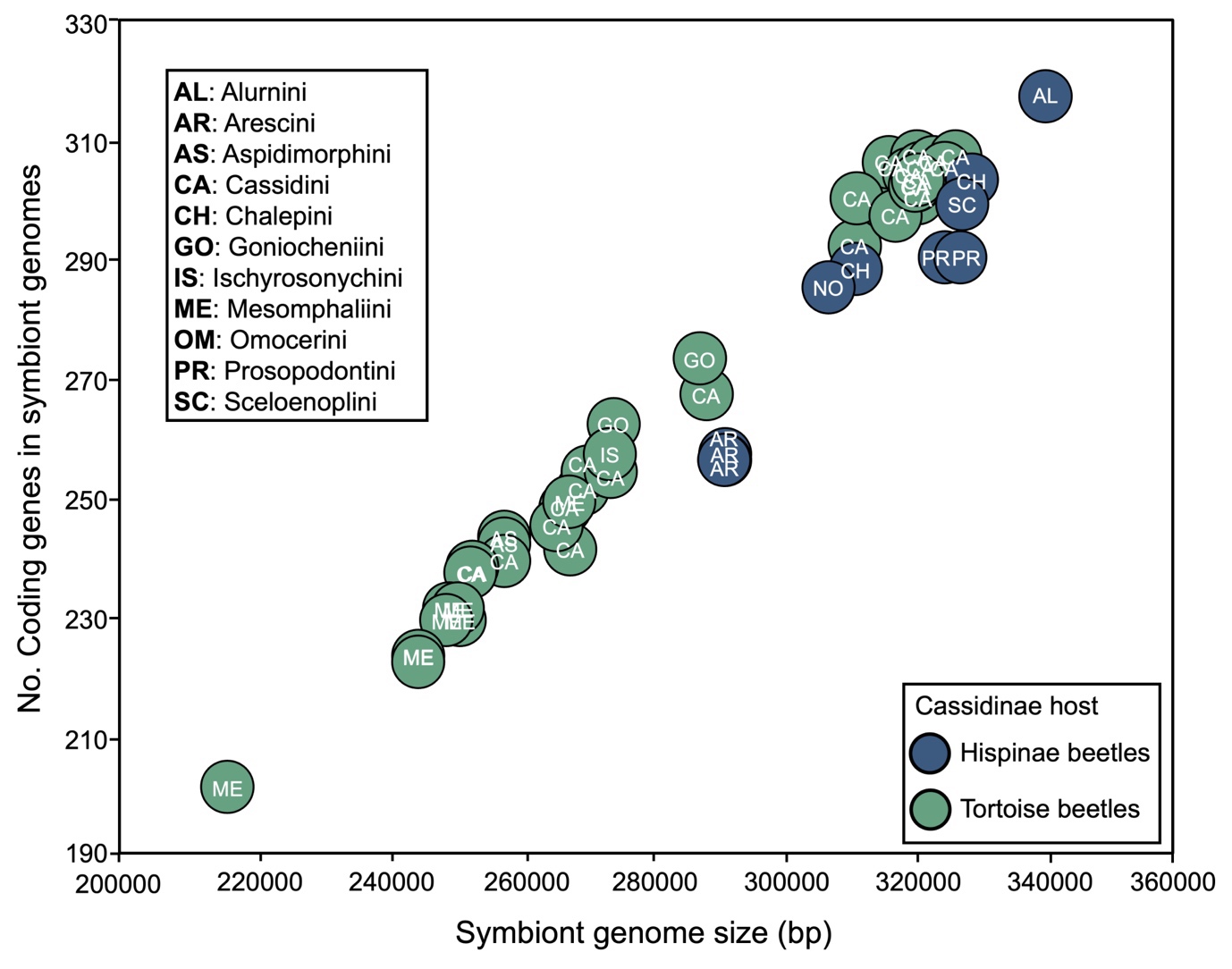

**Figure S1. Number of protein-coding genes in *Stammera* lineages from hispine and tortoise beetles is positively correlated with symbiont genome size** (Spearman’s rank correlation, rho = 0.934, *p* < 0.001). Individual symbiont genomes are labelled with initials of the host tribe.

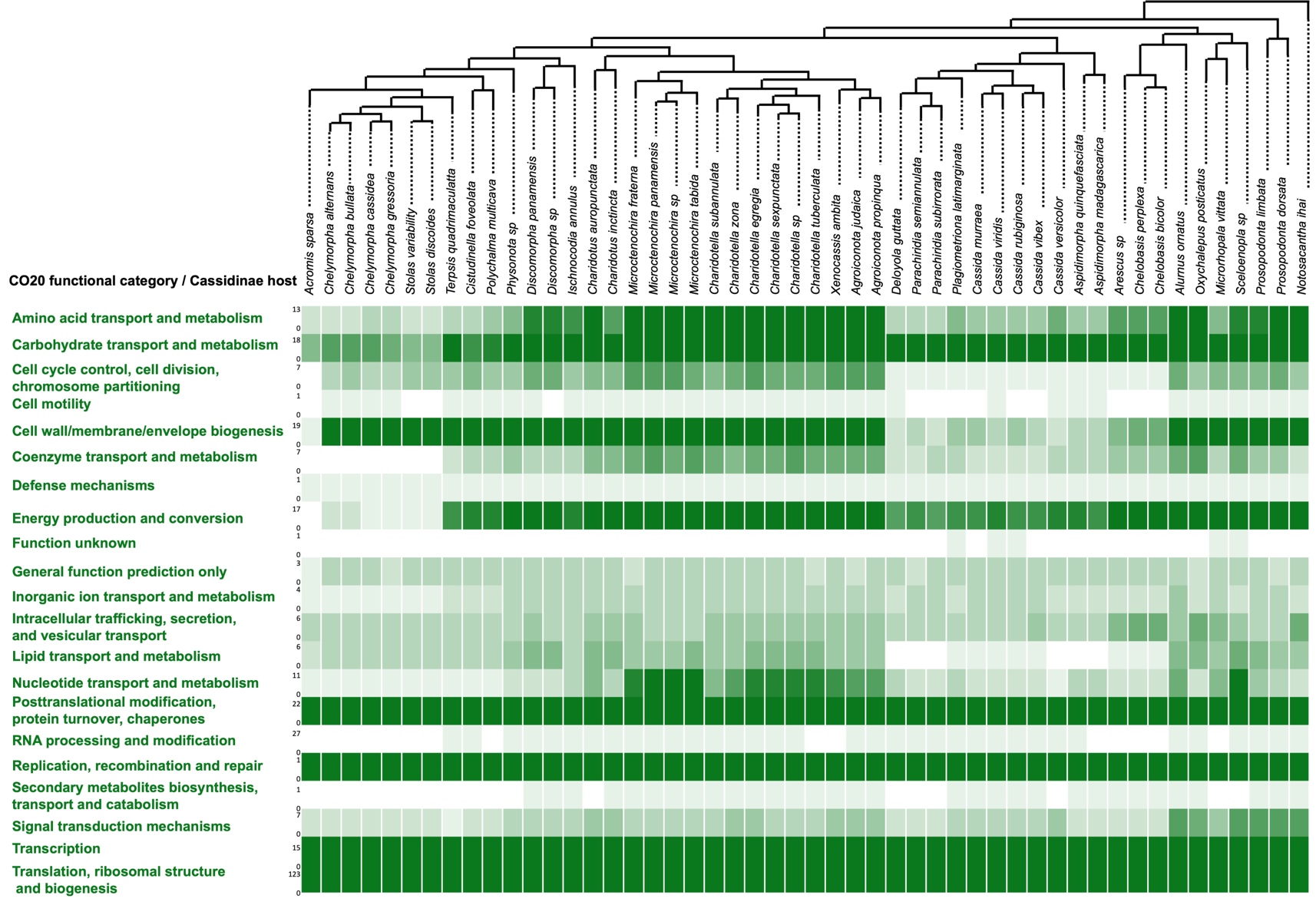

**Figure S2. Frequency of Cluster of Orthologous Genes (COG) functional categories across *Stammera* genomes.**

This figure is based on the pangenome obtained comparing gene functions across genomes rather than sequences. Frequency is displayed by an increasing gradient of colors where grey indicates absence of genes belonging to a functional category and dark green indicates the highest gene frequency of a category in a particular *Stammera* genome. The number at the right of the category name indicates the minimum (bottom) and maximum (top) number of genes belonging to that category.

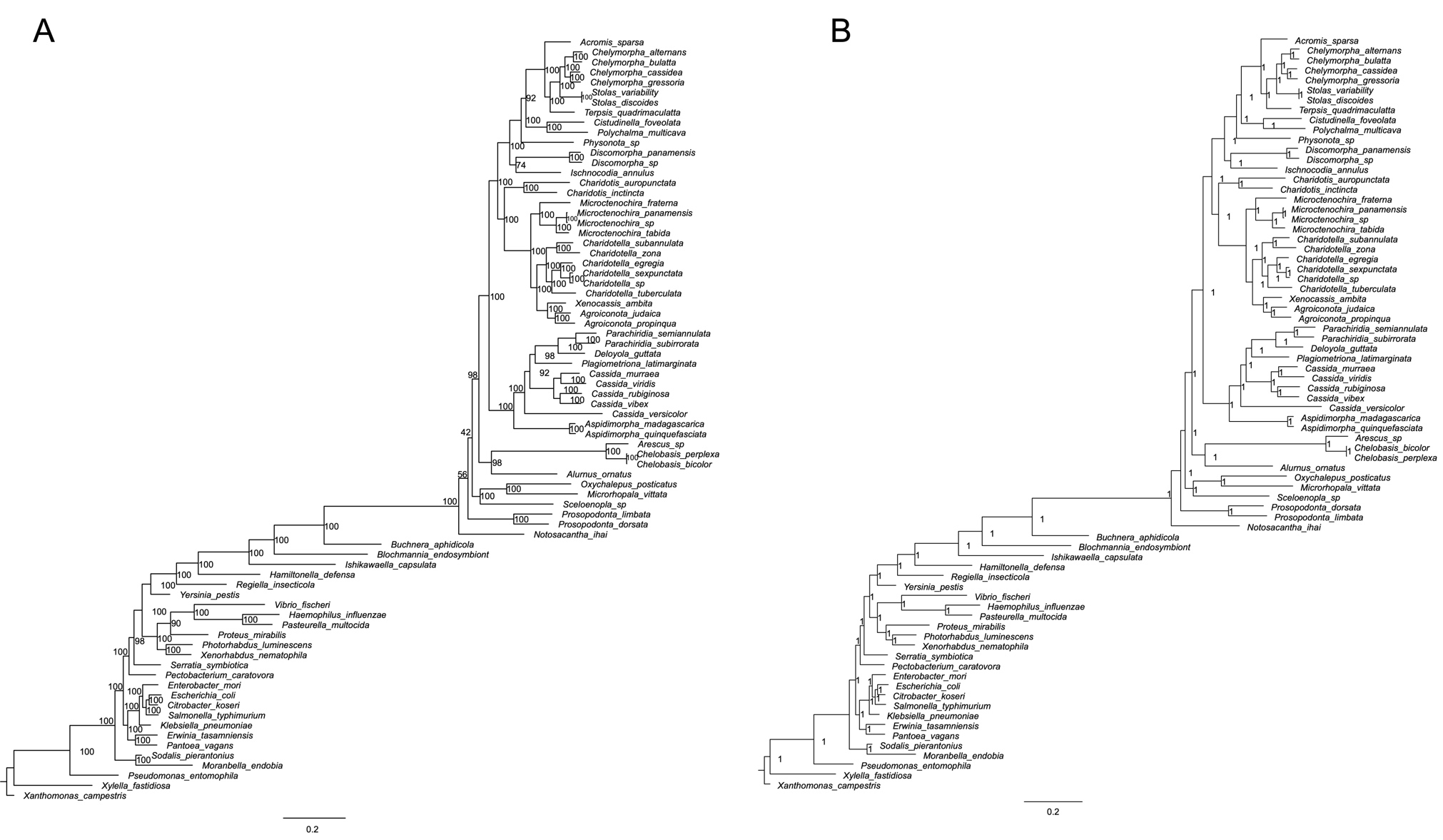

**Figure S3. Full detailed *Stammera* phylogenies including outgroups based on Bayesian (A) and Maximum Likelihood methods (B).** Phylogenomic trees were constructed based on a concatenated alignment of 61 single-copy core genes in MrBayes and RAxML, respectively, using the most appropriate substitution model according to PartitionFinder2. Bayesian posterior probabilities and bootstrap suport values are shown for each node. Bacterial genomes used as outgroups were obtained from NCBI and accession numbers are indicated in Table S5.

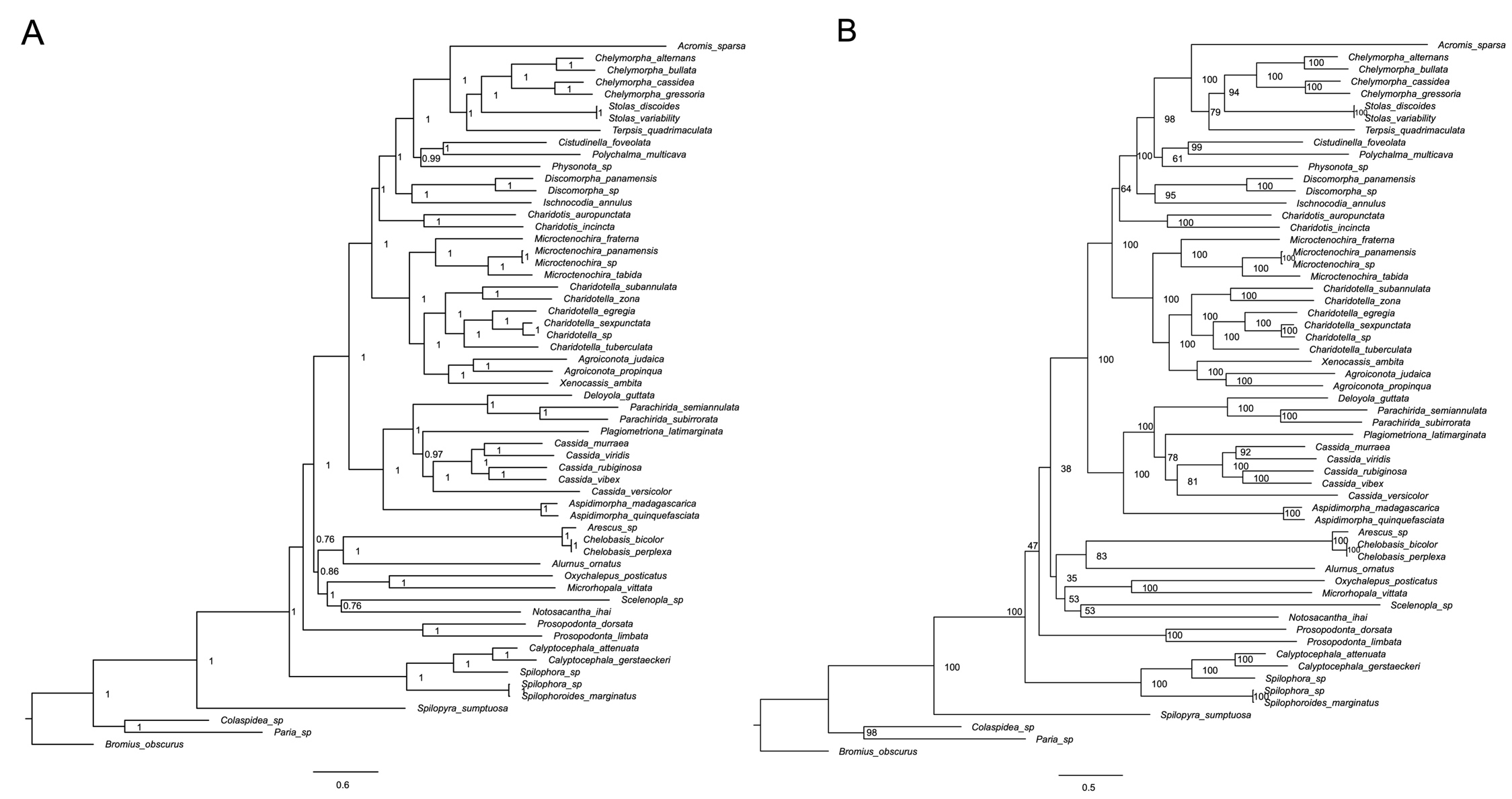

**Figure S4. Full detailed host phylogenies based on 15 mitochondrial genes.** Phylogenetic trees were constructed by Bayesian (A) and Maximum Likelihood (B) methods. Bayesian posterior probabilities and bootstrap suport values are shown for each node.

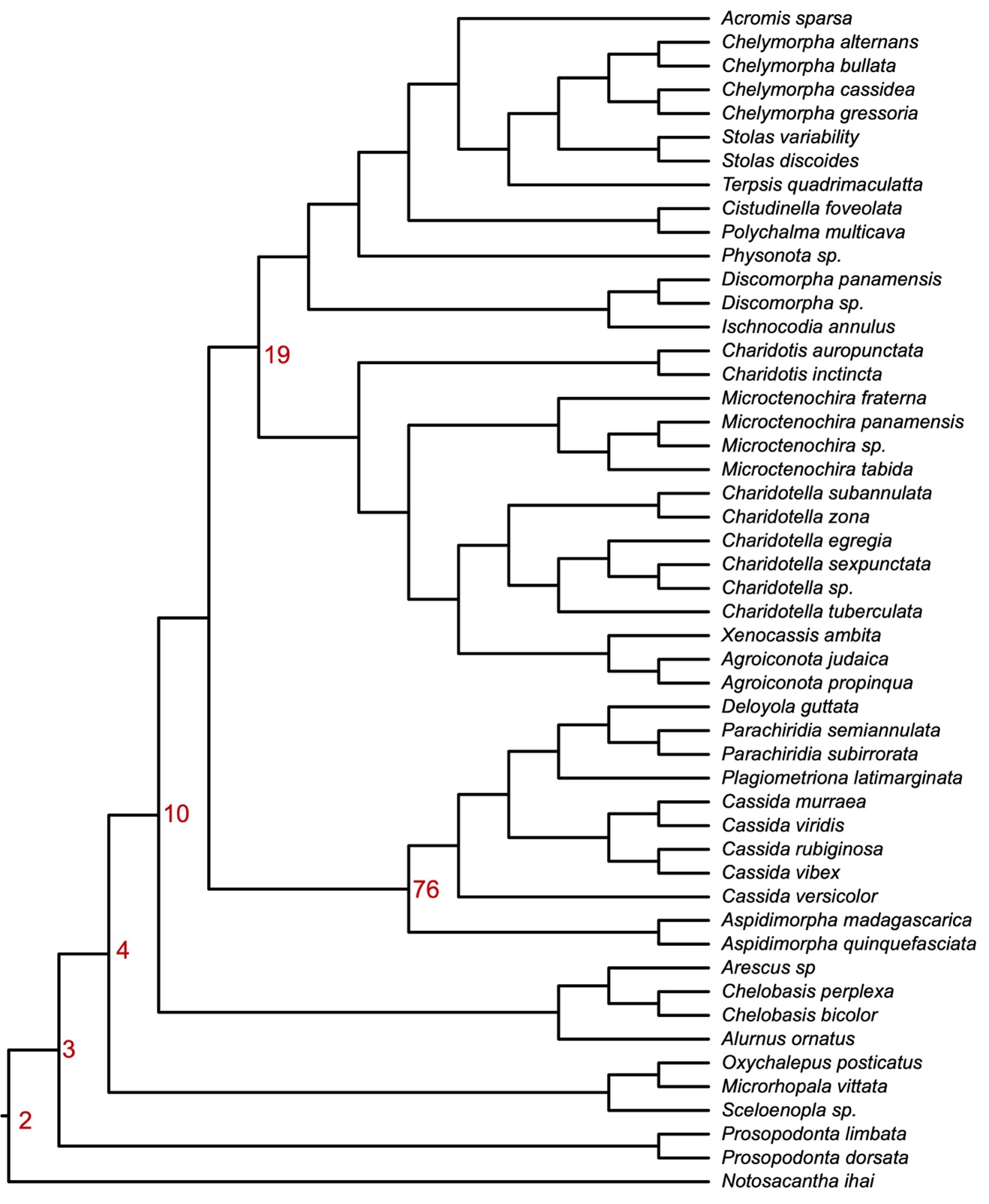

**Figure S5. *Stammera* phylogenomic tree used to infer the ancestral nodes for symbiont genes encoding digestive enzymes.** Tree was constructed based on a concatenated alignment of 124 single-copy *Stammera* core genes. Node numbers are indicated to reference the posterior probabilities for each symbiont gene. Related to Figure 4 and Table S3.

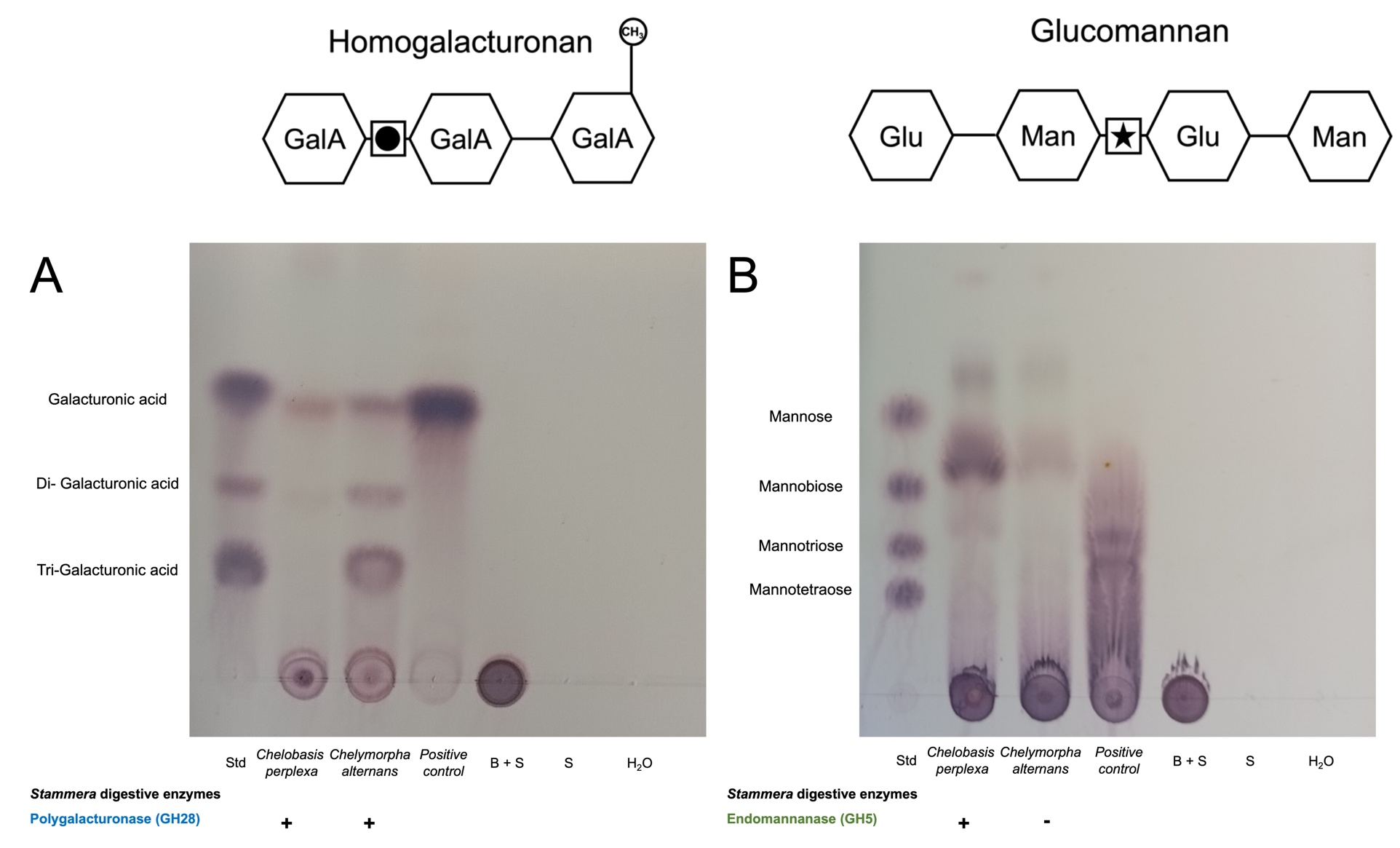

**Figure S6. *Stammera* upgraded the digestive capacity of the hispine beetle *Chelobasis bicolor*.** Thin-layer chromatogram (TLC) illustrating the breakdown products of polygalacturonase activity (A) and endomannanase activity (B) against polygalacturonic acid and glucomannan, respectively. Gut contents from the hispine beetle *Chelobasis bicolor* and the tortoise beetle *Chelymorpha alternans* were used. Polygalacturonase and mannanase from *Aspergillus niger* were used as positive controls. Abbreviations: B, buffer; S, substrate; Std, standard; GalA, galacturonic acid; Glu, glucose; Man, mannose.

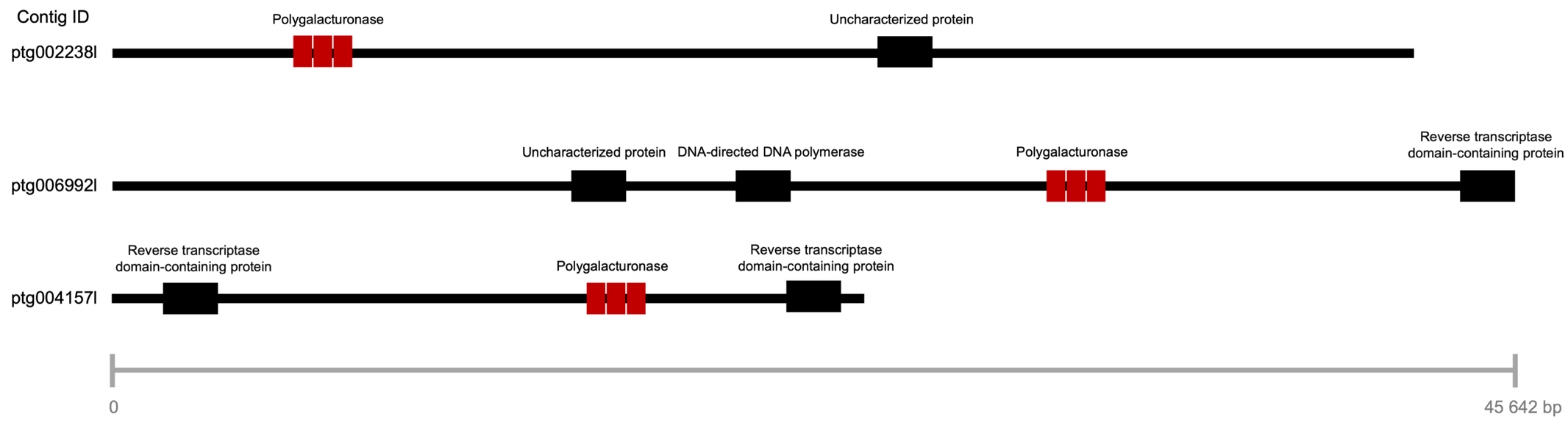

**Figure S7. Schematic representation of polygalacturonase-containing contigs in *Calyptocephala atennuata* draft assembly.** Three copies of the polygalacturonase-encoding gene were identified in *C. attenuata* genome draft assembly generated by PacBio sequencing. Gene structure was conserved across all copies and all of them were flanked by other insect genes based on blast matches.

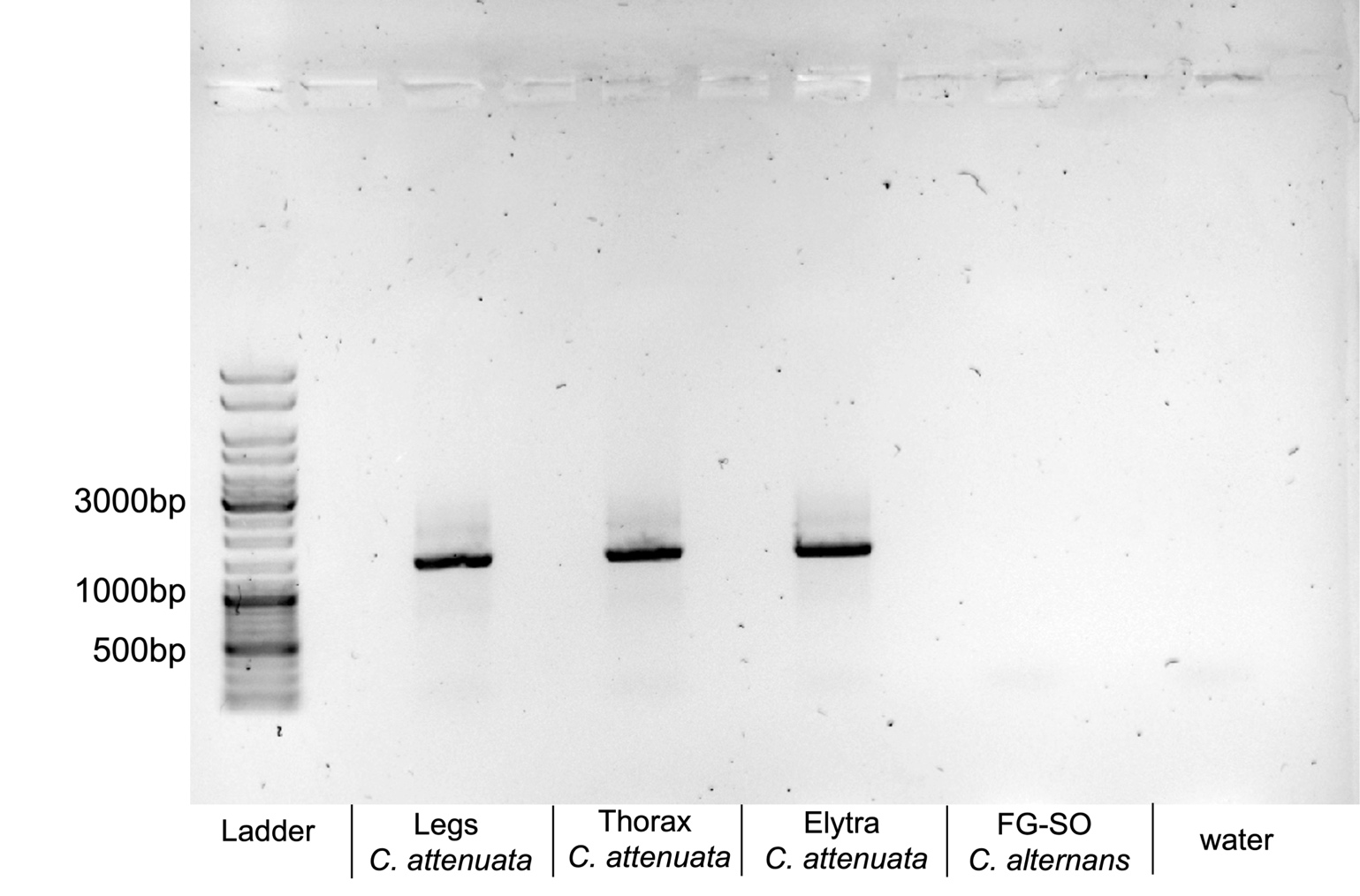

**Figure S8. Polygalacturonase gene identified in *Calyptocephala attenuata* draft assembly is host-encoded.** Electrophoresis gel image displaying the PCR amplification of the host-encoded polygalacturonase (1408 bp fragment size; primers in Table S7) in legs, thorax, and elytra samples from *Calyptocephala attenuata* in rows 2,3, and 4, respectively. A sample from foregut-symbiotic organs (FG-SO) of *Chelymorpha alternans* was also included as a negative control (row 4) in addition to water (row 5). GeneRuler DNA Ladder Mix from Thermo Fisher Scientific was used as a ladder.

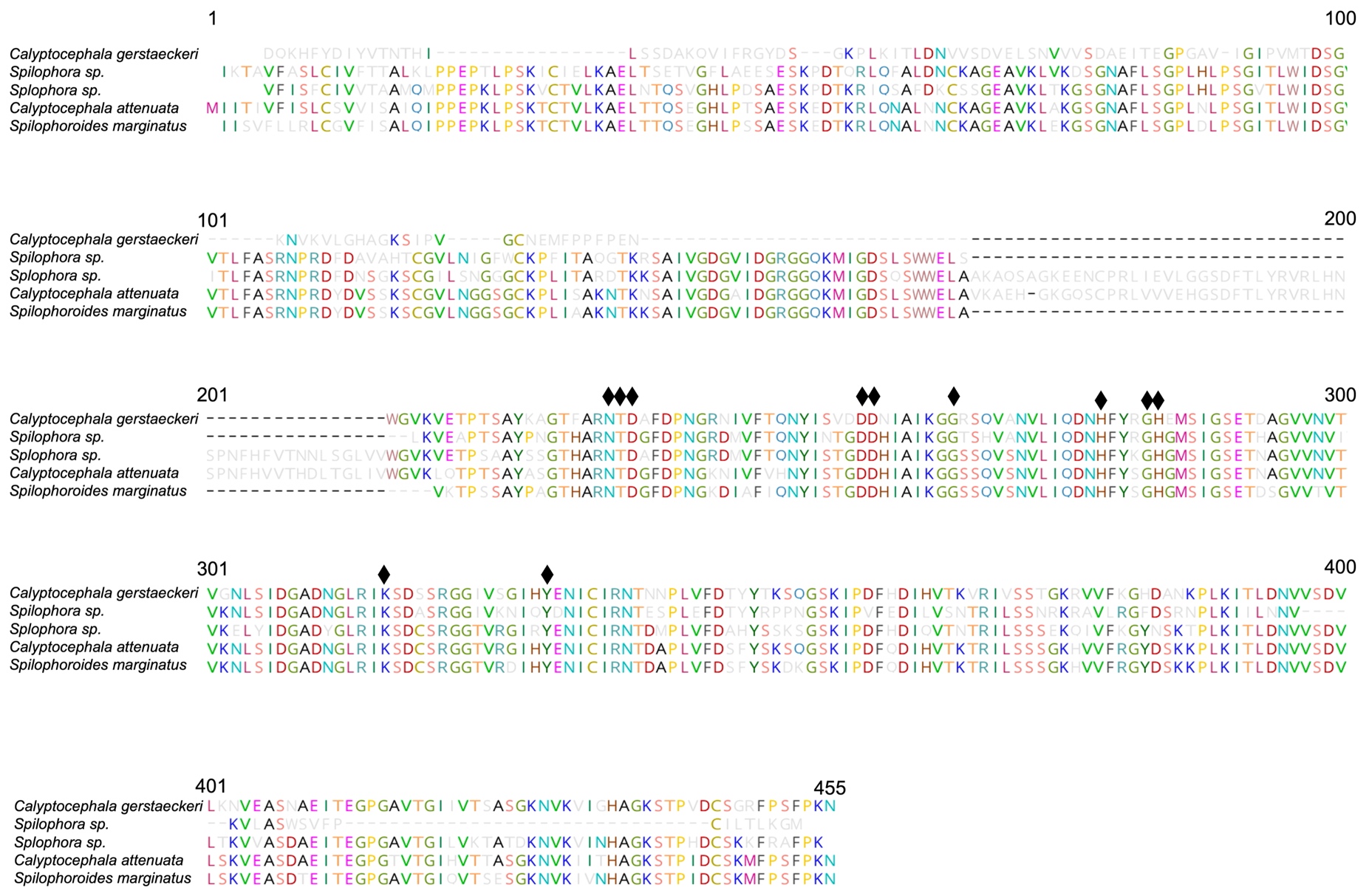

**Figure S9. Host-encoded polygalacturonase gene is present in all members of the Spilophorini.** Polygalacturonase-encoding genes were retrived from metagenomic assemblies for C. *gerstaeckeri, Spilophora sp., Spilophora sp.,* and *Spilophoroides marginatus* based on blast matches of the identified gene in *C. attenuata* by PacBio sequencing. Functionally important amino acids are conserved across all Spilophorini species which are indicated with a diamond.

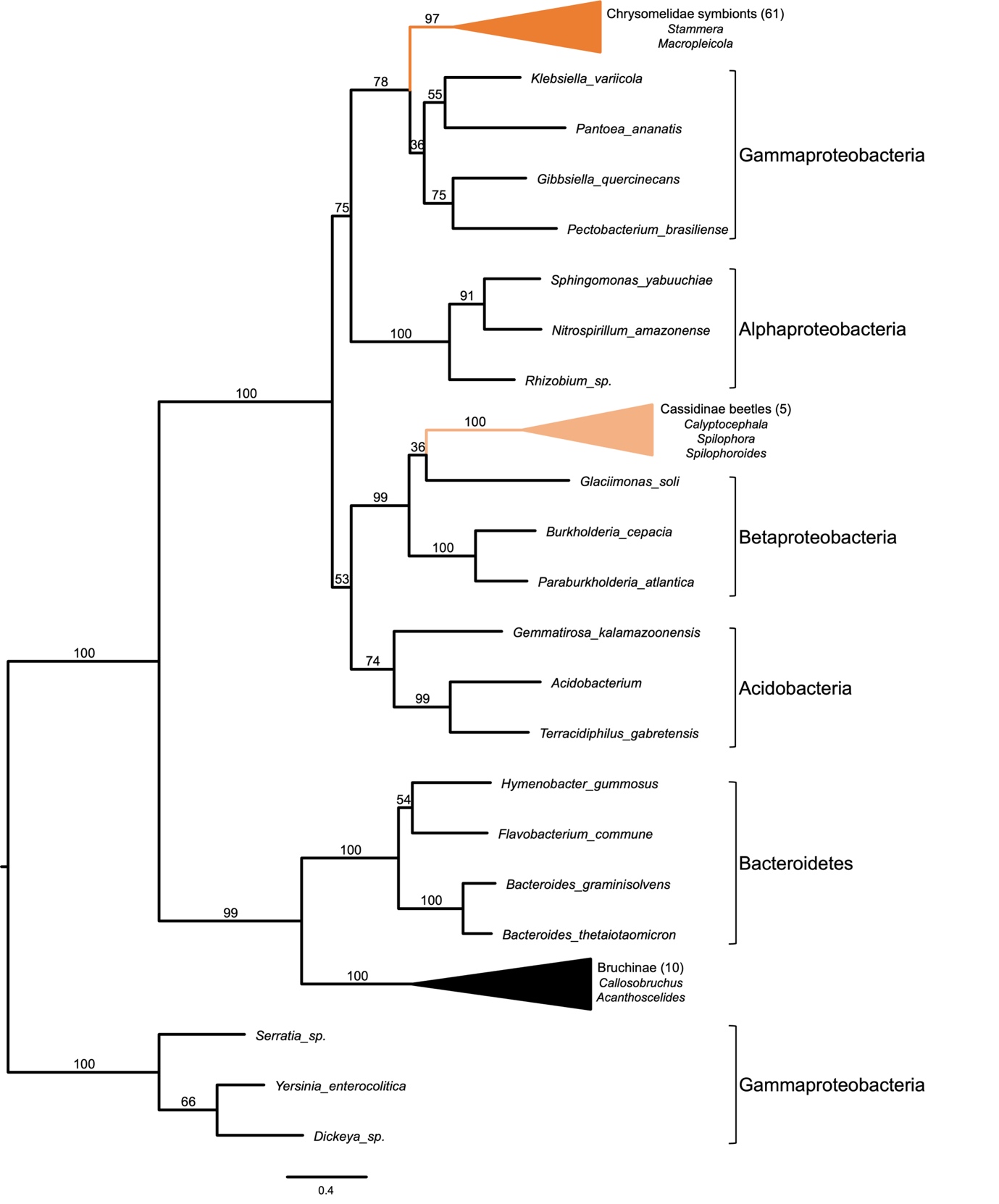

**Figure S10. Phylogenetic placement of Cassidinae encoded GH28 pectinases compared to other Chrysomelidae encoded and symbiont encoded GH28s.** Maximum Likelihood phylogeny is based on an alignment of protein sequences from Proteobacteria, Bacteroidetes, Bruchinae beetles (black triangle), Chrysomelidae symbionts (dark orange triangle) and non-symbiotic Cassidinae (light orange triangle). Triangles indicate collapsed clades.

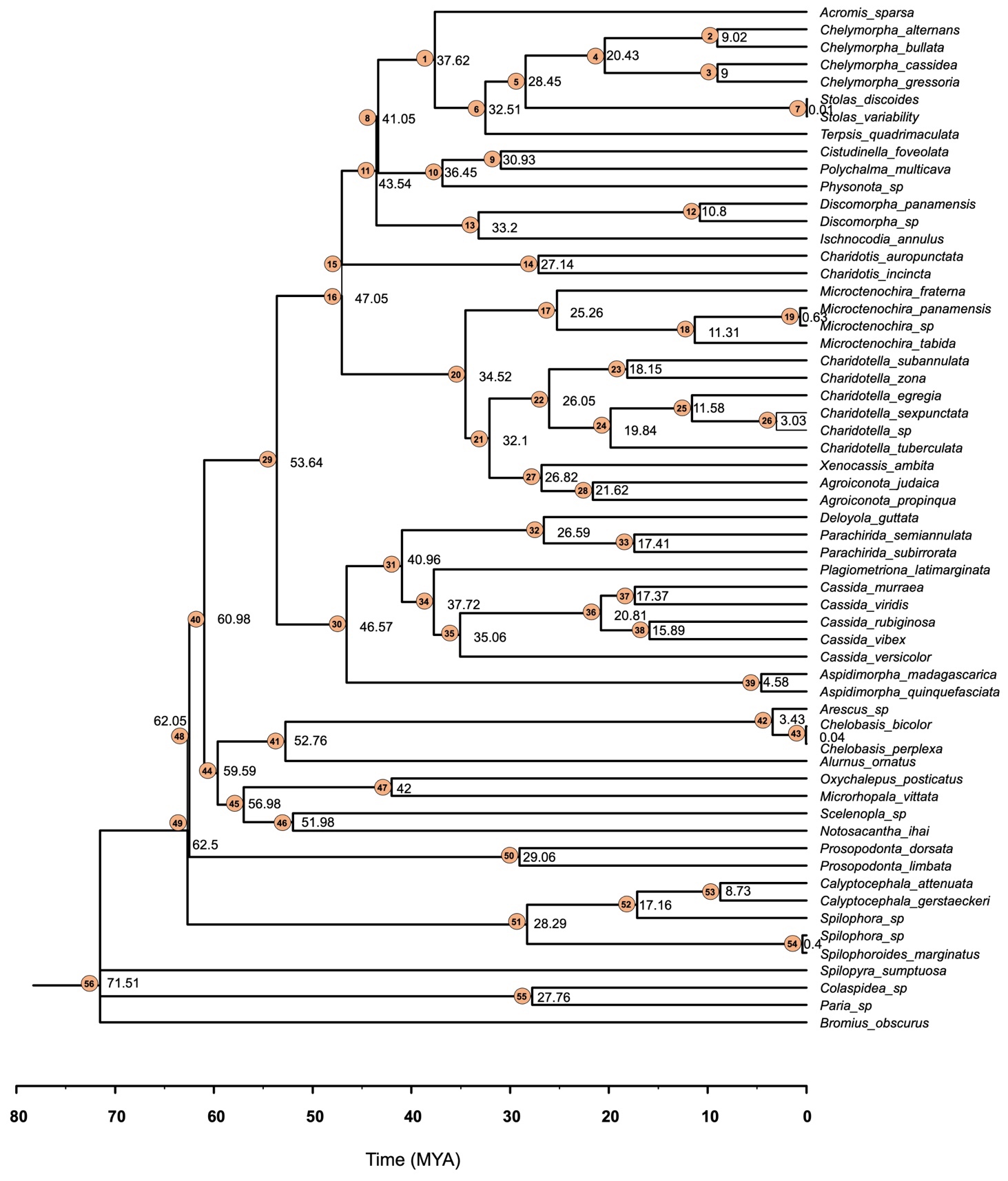

**Figure S11. Divergence-date estimates for the Cassidinae based on 15 mitochondrial genes in BEAST analyses.** Median date estimates (in million years) are indicated for each node. Intervals for each estimate date are included in Table S4.

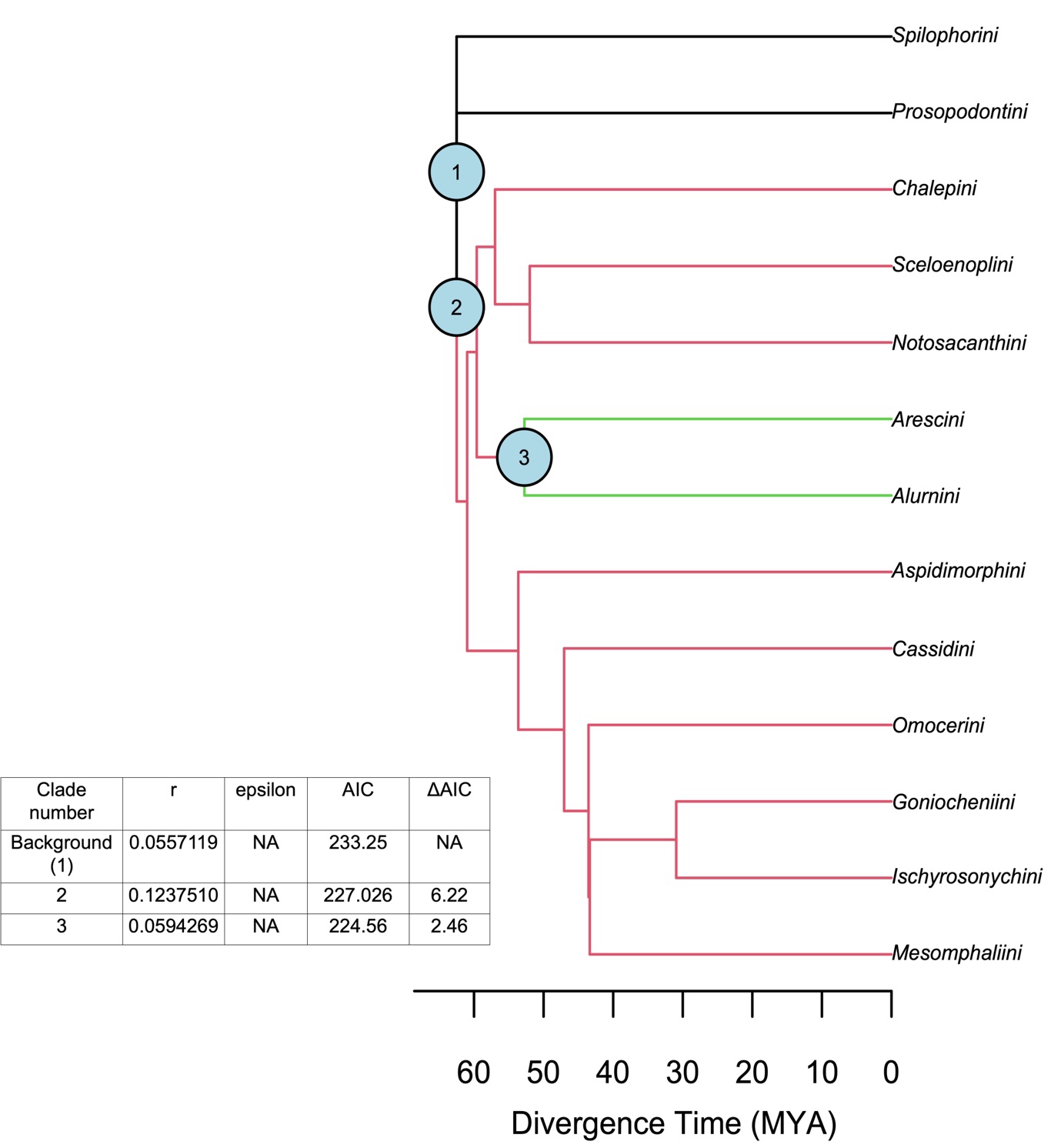

**Figure S12. Tribe-level Cassidinae tree indicating the diversification shifts identified by MEDUSA.** Branch color indicates an increase (red) or a decrease (green) in diversification across the Cassidinae phylogeny. Clade number included in the table represents the rate shifts identified in the tree. Average diversification rate is indicated by r. AIC and ΔAIC show improvement of AIC score over a yule model.

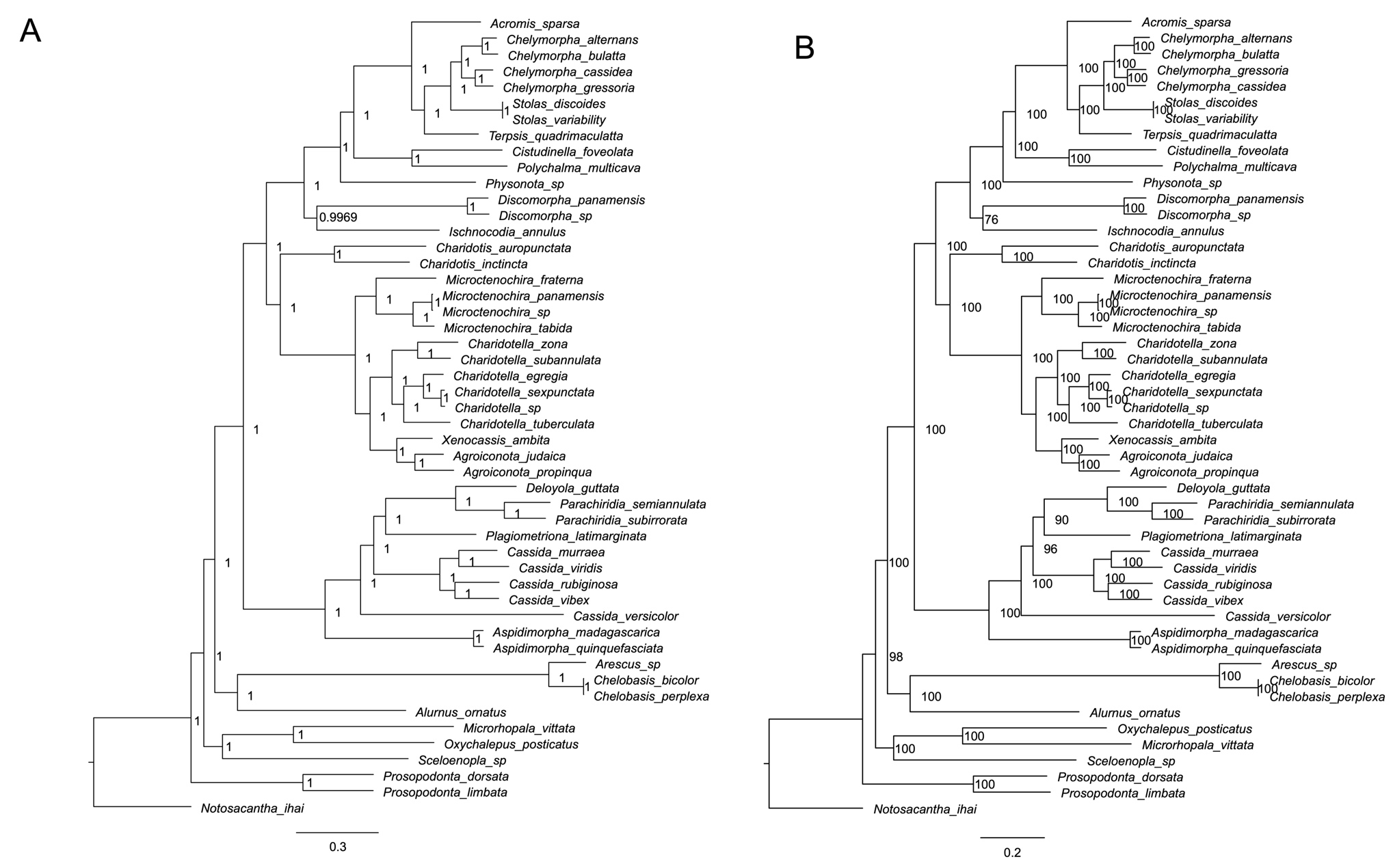

**Figure S13. Unrooted *Stammera* phylogenies based on 124 single-copy core genes constructed by Bayesian (A) and Maximum Likelihood methods (B).** Bayesian posterior probabilities and bootstrap suport values are shown for each node.

| **Beetle species** | **Tribe** | **Location** |
| --- | --- | --- |
| *Alurnus ornatus* | Alurnini | Panama |
| *Arescus sp.* | Arescini | Panama |
| *Chelobasis bicolor* | Arescini | Panama |
| *Chelobasis perplexa* | Arescini | Panama |
| *Aspidimorpha madagascarica* | Aspidimorphini | La Reunion, France |
| *Aspidimorpha quinquefasciata* | Aspidimorphini | La Reunion, France |
| *Agroiconota judaica* | Cassidini | Panama |
| *Agroiconota propinqua* | Cassidini | Panama |
| *Cassida murraea* | Cassidini | Germany |
| *Cassida rubiginosa* | Cassidini | New Zealand |
| *Cassida versicolor* | Cassidini | Japan |
| *Cassida vibex* | Cassidini | Germany |
| *Cassida viridis* | Cassidini | Germany |
| *Charidotella egregia* | Cassidini | Panama |
| *Charidotella sexpunctata* | Cassidini | USA |
| *Charidotella sp.* | Cassidini | Martinique, France |
| *Charidotella subannulata* | Cassidini | Panama |
| *Charidotella tuberculata* | Cassidini | Panama |
| *Charidotella zona* | Cassidini | Panama |
| *Charidotus auropunctata* | Cassidini | Panama |
| *Charidotus inctincta* | Cassidini | Panama |
| *Deloyola guttata* | Cassidini | Panama |
| *Ischnocodia annulus* | Cassidini | Panama |
| *Microctenochira fraterna* | Cassidini | Panama |
| *Microctenochira panamensis* | Cassidini | Panama |
| *Microctenochira sp.* | Cassidini | Panama |
| *Microctenochira tabida* | Cassidini | Panama |
| *Parachiridia semiannulata* | Cassidini | Panama |
| *Parachiridia subirrorata* | Cassidini | Panama |
| *Plagiometriona latimarginata* | Cassidini | Panama |
| *Xenocassis ambita* | Cassidini | Panama |
| *Microrhopala vittata* | Cassidini | Panama |
| *Oxychalepus posticatus* | Cassidini | Panama |
| *Physonota sp.* | Goniocheniini | Panama |
| *Polychalma multicava* | Goniocheniini | Panama |
| *Cistudinella foveolata* | Ischyrosonychini | Panama |
| *Acromis sparsa* | Mesomphaliini | Panama |
| *Chelymorpha alternans* | Mesomphaliini | Germany |
| *Chelymorpha bullata* | Mesomphaliini | Panama |
| *Chelymorpha cassidea* | Mesomphaliini | USA |
| *Chelymorpha gressoria* | Mesomphaliini | Panama |
| *Stolas discoides* | Mesomphaliini | Panama |
| *Stolas variability* | Mesomphaliini | Panama |
| *Terpsis quadrimaculatta* | Mesomphaliini | Panama |
| *Notosacantha ihai* | Notosocanthini | Japan |
| *Discomorpha panamensis* | Omocerini | Panama |
| *Discomorpha sp.* | Omocerini | Panama |
| *Prosopodonta dorsata* | Prosopodontini | Panama |
| *Prosopodonta limbata* | Prosopodontini | Panama |
| *Sceloenopla sp.* | Sceloenoplini | Panama |
| *Calyptocephala attenuata* | Spilophorini | Panama |
| *Calyptocephala gerstaeckeri* | Spilophorini | Panama |
| *Spilophoroides marginatus* | Spilophorini | Panama |
| *Spilophora sp.* | Spilophorini | Panama |
| *Spilophora sp.* | Spilophorini | Panama |

**Table S1. List of Cassidinae species included in this study, tribe and country of collection.**

**Table S2. Genomic features of *Stammera* included in this study.**

| **Species** | **Tribe** | **Group** | **Genome size** | **Plasmids** | **AT Content (%)** | **CDS** | **rRNA** | **tRNA** | **tmRNA** | **misc RNA** | **Pseudogenes** |
| --- | --- | --- | --- | --- | --- | --- | --- | --- | --- | --- | --- |
| *Alurnus ornatus* | Alurnini | Hispine beetles | 340730 | 2 | 18.4 | 317 | 3 | 31 | 1 | 2 | 6 |
| *Arescus sp.* | Arescini | Hispine beetles | 291998 | 2 | 20 | 257 | 3 | 30 | 1 | 2 | 6 |
| *Chelobasis bicolor* | Arescini | Hispine beetles | 291995 | 2 | 19.6 | 256 | 3 | 30 | 1 | 3 | 7 |
| *Chelobasis perplexa* | Arescini | Hispine beetles | 291902 | 2 | 19.6 | 256 | 3 | 30 | 1 | 3 | 7 |
| *Aspidimorpha madagascarica* | Aspidimorphini | Tortoise beetles | 258484 | 1 | 16.5 | 243 | 3 | 29 | 1 | 3 | 2 |
| *Aspidimorpha quinquefasciata* | Aspidimorphini | Tortoise beetles | 258530 | 1 | 16.5 | 242 | 3 | 29 | 1 | 2 | 2 |
| *Agroiconota judaica* | Cassidini | Tortoise beetles | 317369 | 2 | 17.4 | 305 | 3 | 29 | 1 | 2 | 0 |
| *Agroiconota propinqua* | Cassidini | Tortoise beetles | 316928 | 2 | 17.6 | 306 | 3 | 29 | 1 | 2 | 1 |
| *Cassida murraea* | Cassidini | Tortoise beetles | 268530 | 2 | 15.1 | 241 | 3 | 29 | 1 | 2 | 9 |
| *Cassida rubiginosa* | Cassidini | Tortoise beetles | 270381 | 2 | 15.4 | 251 | 3 | 29 | 1 | 2 | 1 |
| *Cassida versicolor* | Cassidini | Tortoise beetles | 267681 | 2 | 16.2 | 248 | 3 | 28 | 1 | 2 | 4 |
| *Cassida vibex* | Cassidini | Tortoise beetles | 271169 | 2 | 15.2 | 254 | 3 | 29 | 1 | 2 | 1 |
| *Cassida viridis* | Cassidini | Tortoise beetles | 266462 | 2 | 16 | 245 | 3 | 29 | 1 | 2 | 5 |
| *Charidotella egregia* | Cassidini | Tortoise beetles | 321123 | 2 | 17.9 | 307 | 3 | 29 | 1 | 2 | 0 |
| *Charidotella sexpunctata* | Cassidini | Tortoise beetles | 319936 | 2 | 18 | 304 | 3 | 29 | 1 | 2 | 2 |
| *Charidotella sp.* | Cassidini | Tortoise beetles | 321733 | 2 | 17.9 | 305 | 3 | 29 | 1 | 2 | 2 |
| *Charidotella subannulata* | Cassidini | Tortoise beetles | 311669 | 2 | 17 | 292 | 3 | 29 | 1 | 2 | 3 |
| *Charidotella tuberculata* | Cassidini | Tortoise beetles | 321341 | 2 | 17.8 | 300 | 3 | 29 | 1 | 2 | 4 |
| *Charidotella zona* | Cassidini | Tortoise beetles | 312082 | 2 | 18.2 | 300 | 5 | 31 | 1 | 2 | 2 |
| *Charidotus auropunctata* | Cassidini | Tortoise beetles | 321044 | 2 | 18.8 | 302 | 3 | 30 | 1 | 3 | 6 |
| *Charidotus inctincta* | Cassidini | Tortoise beetles | 317910 | 2 | 16.9 | 297 | 3 | 30 | 1 | 2 | 1 |
| *Deloyola guttata* | Cassidini | Tortoise beetles | 258495 | 2 | 15 | 239 | 3 | 29 | 1 | 2 | 6 |
| *Ischnocodia annulus* | Cassidini | Tortoise beetles | 289195 | 1 | 15.1 | 267 | 3 | 29 | 1 | 2 | 9 |
| *Microctenochira fraterna* | Cassidini | Tortoise beetles | 320874 | 2 | 17.5 | 302 | 3 | 29 | 1 | 2 | 4 |
| *Microctenochira panamensis* | Cassidini | Tortoise beetles | 323662 | 2 | 17.3 | 306 | 3 | 29 | 1 | 2 | 0 |
| *Microctenochira sp.* | Cassidini | Tortoise beetles | 327024 | 2 | 17.2 | 307 | 3 | 29 | 1 | 2 | 2 |
| *Microctenochira tabida* | Cassidini | Tortoise beetles | 325345 | 2 | 17.9 | 305 | 3 | 29 | 1 | 3 | 2 |
| *Parachiridia semiannulata* | Cassidini | Tortoise beetles | 253694 | 2 | 15.2 | 238 | 3 | 29 | 1 | 3 | 2 |
| *Parachiridia subirrorata* | Cassidini | Tortoise beetles | 253400 | 2 | 15.1 | 237 | 3 | 29 | 1 | 3 | 2 |
| *Plagiometriona latimarginata* | Cassidini | Tortoise beetles | 274660 | 2 | 14.8 | 254 | 3 | 29 | 1 | 3 | 1 |
| *Xenocassis ambita* | Cassidini | Tortoise beetles | 321242 | 2 | 17 | 303 | 3 | 29 | 1 | 3 | 4 |
| *Microrhopala vittata* | Chalepini | Hispine beetles | 311863 | 2 | 20.4 | 288 | 3 | 31 | 1 | 2 | 2 |
| *Oxychalepus posticatus* | Chalepini | Hispine beetles | 329508 | 2 | 19 | 303 | 3 | 30 | 1 | 2 | 4 |
| *Physonota sp.* | Goniocheniini | Tortoise beetles | 288173 | 1 | 16.1 | 273 | 3 | 30 | 1 | 2 | 1 |
| *Polychalma multicava* | Goniocheniini | Tortoise beetles | 275070 | 1 | 16 | 262 | 3 | 29 | 1 | 2 | 5 |
| *Cistudinella foveolata* | Ischyrosonychini | Tortoise beetles | 274528 | 1 | 15.4 | 257 | 3 | 29 | 1 | 2 | 2 |
| *Acromis sparsa* | Mesomphaliini | Tortoise beetles | 216430 | 1 | 13.9 | 201 | 3 | 29 | 1 | 2 | 0 |
| *Chelymorpha alternans* | Mesomphaliini | Tortoise beetles | 250099 | 1 | 15.5 | 231 | 3 | 29 | 1 | 2 | 4 |
| *Chelymorpha bullata* | Mesomphaliini | Tortoise beetles | 251774 | 1 | 15.3 | 229 | 3 | 29 | 1 | 2 | 4 |
| *Chelymorpha cassidea* | Mesomphaliini | Tortoise beetles | 251492 | 1 | 15.3 | 231 | 3 | 29 | 1 | 2 | 3 |
| *Chelymorpha gressoria* | Mesomphaliini | Tortoise beetles | 249733 | 1 | 15.4 | 229 | 3 | 29 | 1 | 3 | 4 |
| *Stolas discoides* | Mesomphaliini | Tortoise beetles | 245391 | 1 | 14.5 | 223 | 3 | 29 | 1 | 2 | 3 |
| *Stolas variability* | Mesomphaliini | Tortoise beetles | 245433 | 1 | 14.5 | 222 | 3 | 29 | 1 | 2 | 4 |
| *Terpsis quadrimaculatta* | Mesomphaliini | Tortoise beetles | 268407 | 0 | 14.8 | 249 | 3 | 29 | 1 | 3 | 3 |
| *Notosacantha ihai* | Notosocanthini | Hispine beetles | 307768 | 2 | 16.9 | 285 | 3 | 30 | 1 | 2 | 13 |
| *Discomorpha panamensis* | Omocerini | Tortoise beetles | 291398 | 1 | 16.6 | 278 | 3 | 29 | 1 | 3 | 2 |
| *Discomorpha sp.* | Omocerini | Tortoise beetles | 290616 | 1 | 16.3 | 274 | 3 | 29 | 1 | 2 | 4 |
| *Prosopodonta dorsata* | Prosopodontini | Hispine beetles | 325465 | 2 | 19 | 290 | 3 | 31 | 1 | 3 | 9 |
| *Prosopodonta limbata* | Prosopodontini | Hispine beetles | 327731 | 2 | 20.1 | 290 | 3 | 31 | 1 | 2 | 8 |
| *Sceloenopla sp.* | Sceloenoplini | Hispine beetles | 328085 | 2 | 17.7 | 299 | 3 | 28 | 1 | 2 | 7 |

**Table S3. Probabilities of the ancestral state reconstructions for the four digestive enzymes identified in *Stammera*’s accessory genome.** Values shown are for the indicated nodes in which a major change, either acquisition or loss of the specified genes occurred. Related to Figure 4 and Figure

S5.

|  | **Posterior probabilities** | | | |
| --- | --- | --- | --- | --- |
| **Node\Character** | **Alpha glucuronidase (GH67)** | **Rhamnogalacturonan lyase**  **(PL4)** | **Endomannanase**  **(GH5)** | **Pectin methylesterase**  **(CE8)** |
| node 2 | 0.999 | 0.947 | 0.00015 | 0.00015 |
| node 3 | 0.999 | 0.941 | 0.000 | 0.000 |
| node 4 | 0.999 | 0.951 | 0.000 | 0.000 |
| node 10 | 0.999 | 0.934 | 0.000 | 0.000 |
| node 13 | 0.999 | 0.006 | 0.998 | 0.998 |
| node 19 | 0.999 | 0.016 | 0.000 | 0.000 |
| node 76 | 0.026 | 0.99 | 0.000 | 0.000 |

**Table S4. Divergence-date estimates (millions of years) for all the nodes across the Cassidinae obtained from BEAST.** Median estimates and the 95% highest posterior density intervals are included. Related to Figure S10.

| **Node** | **Median estimate** | **95% Confidence interval** |
| --- | --- | --- |
| 1 | 37.62 | 35.43 - 40 |
| 2 | 9.02 | 8.26 - 9.77 |
| 3 | 9 | 8.26 – 9.76 |
| 4 | 20.43 | 19.02 – 21.77 |
| 5 | 28.45 | 26.63 – 30.21 |
| 6 | 32.51 | 30.5 – 34.49 |
| 7 | 0.01 | 0 – 0.02 |
| 8 | 41.05 | 38.95 – 43.43 |
| 9 | 30.93 | 28.71 – 33.23 |
| 10 | 36.45 | 34.03 – 38.67 |
| 11 | 43.54 | 41.19 – 45.82 |
| 12 | 10.8 | 9.91 – 11.67 |
| 13 | 33.2 | 30.7 – 35.69 |
| 14 | 27.14 | 25.15 – 29.22 |
| 16 | 47.05 | 44.64 – 49.57 |
| 17 | 25.26 | 23.55 – 27 |
| 18 | 11.31 | 10.45 – 12.24 |
| 19 | 0.63 | 0.53 – 0.74 |
| 20 | 34.52 | 32.56 – 36.4 |
| 21 | 32.1 | 30.27 – 33.88 |
| 22 | 26.05 | 24.39 – 27.59 |
| 23 | 18.15 | 16.75 – 19.45 |
| 24 | 19.84 | 18.49 – 21.23 |
| 25 | 11.58 | 10.68 – 12.46 |
| 26 | 3.03 | 2.73 – 3.35 |
| 27 | 26.82 | 25.23 – 28.62 |
| 28 | 21.62 | 20.12 – 23.16 |
| 29 | 53.64 | 50.97 – 56.42 |
| 30 | 46.57 | 44.11 – 49.08 |
| 31 | 40.96 | 38.83 – 43.1 |
| 32 | 26.59 | 24.89 – 28.36 |
| 33 | 17.41 | 16.16 – 18.72 |
| 34 | 37.72 | 35.61 – 39.79 |
| 35 | 35.06 | 33.19 – 36.96 |
| 36 | 20.81 | 19.53 – 22.12 |
| 37 | 17.37 | 16.14 – 18.68 |
| 38 | 15.89 | 14.68 – 17.07 |
| 39 | 4.58 | 4.14 – 5.06 |
| 40 | 60.98 | 58.06 – 63.96 |
| 41 | 52.76 | 49.68 – 56.27 |
| 42 | 3.43 | 3.07 – 3.74 |
| 43 | 0.04 | 0.02 – 0.06 |
| 44 | 59.59 | 56.91 – 62.65 |
| 45 | 56.98 | 54.23 – 59.75 |
| 46 | 51.98 | 49.68 – 54.33 |
| 47 | 42 | 39.8 – 44.13 |
| 48 | 62.05 | 59.99 – 64.34 |
| 49 | 62.5 | 59.37 – 65.6 |
| 50 | 29.06 | 26.84 – 31.11 |
| 51 | 28.29 | 26.37 – 30.23 |
| 52 | 17.16 | 15.96 – 18.43 |
| 53 | 8.73 | 8 – 9.47 |
| 54 | 0.4 | 0.31 – 0.48 |
| 55 | 27.76 | 25.87 – 29.83 |
| 56 | 71.56 | 67.6 – 75.36 |

**Table S5. Reference Sequence Database (RefSeq) accession numbers for bacterial outgroups utilized to construct symbiont phylogenies in Figure S3.**

| **Species** | **RefSeq** |
| --- | --- |
| Blochmannia endosymbiont of Camponotus modoc | GCF_023585785.1 |
| *Buchnera aphidicola* | GCF_003099975.1 |
| *Candidatus* Hamiltonella defensa | GCF_000021705.1 |
| *Candidatus* Ishikawaella capsulata Mpkobe | GCF_000828515.1 |
| *Candidatus* Moranella endobia | GCF_000364725.1 |
| *Candidatus* Regiella insecticola | GCF_013373955.1 |
| *Candidatus* Sodalis pierantonius str. SOPE | GCF_000517405.1 |
| *Citrobacter koseri* | GCF_000018045.1 |
| *Enterobacter mori* | GCF_022014715.1 |
| *Erwinia tasamniensis* | GCF_000026185.1 |
| *Escherichia coli* | GCF_000005845.2 |
| *Haemophilus influenzae* | GCF_000931575.1 |
| *Klebsiella pneumoniae* | GCF_000240185.2 |
| *Pantoea vagans* | GCF_004792415.1 |
| *Pasteurella multocida* | GCF_002073255.2 |
| *Pectobacterium caratovora* | GCF_013488025.1 |
| *Photorhabdus luminescens* | GCF_001083805.1 |
| *Proteus mirabilis* | GCF_000069965.1 |
| *Pseudomonas entomophila* | GCF_000026105.1 |
| *Salmonella typhimurium* | GCF_000006945.2 |
| *Serratia symbiotica* | GCF_009831665.3 |
| *Vibrio fischeri* | GCF_000020845.1 |
| *Xanthomonas campestris* | GCF_013388375.1 |
| *Xenorhabdus nematophila* | GCF_014295015.1 |
| *Xylella fastidiosa* | GCF_000007245.1 |
| *Yersinia pestis* | GCF_024498375.1 |

**Table S6. Partitions schemes and the best-fitting models for host and symbiont alignments identified by PartitionFinder2.** Related to Figure S3, S4 and S12.

| **Alignment** | **Subset** | **Partitions by nucleotide position** | **Best model** |
| --- | --- | --- | --- |
| **Host** | 1 | 7550-7800\3, 7801-9402\3, 6339-7548\3, 4422-5295\3, 5298-6145\3, 6148-6337\3, 1804-3305\3, 754-1802\3, 3306-3869\3, 1-562\3, 3871-4421\3 | GTR+I+G |
|  | 2 | 7802-9402\3, 7551-7800\3, 2-562\3 | GTR+I+G |
|  | 3 | 1803-3305\3, 3870-4421\3, 3308-3869\3, 753-1802\3, 3-562\3 | GTR+I+G |
|  | 4 | 9404-9893\3, 563-752\3, 9403-9893\3, 5297-6145\3, 6147-6337\3, 564-752\3, 5296-6145\3, 9405-9893\3 | GTR+I+G |
|  | 5 | 565-752\3, 6146-6337\3 | GTR+I+G |
|  | 6 | 755-1802\3 | GTR+G |
|  | 7 | 1805-3305\3 | GTR+G |
|  | 8 | 3307-3869\3 | GTR+I+G |
|  | 9 | 3872-4421\3 | GTR+I+G |
|  | 10 | 4423-5295\3 | GTR+I+G |
|  | 11 | 11106-11846 9894-11105 4424-5295\3 6338-7548\3 7803-9402\3 7549-7800\3 | GTR+G |
|  | 12 | 6340-7548\3 | GTR+I+G |
| **Rooted phylogeny** | | | |
| **Symbiont** | 1 | 34897-36425\3, 64602-65746\3, 6255-6946\3, 27884-29817\3, 60176-60969\3, 31237-31695\3, 49312-49914\3, 19699-20628\3, 43363-43956\3, 51455-51855\3, 7377-7852\3, 43958-44316\3, 4365-4858\3, 27095-27882\3, 36427-37097\3, 22461-22768\3, 6947-7375\3, 20629-22459\3, 37098-37778\3, 14651-15067\3, 27883-29817\3, 7376-7852\3, 10007-14323\3, 55768-60174\3, 2704-3614\3, 616-909\3, 31696-32568\3, 2086-2259\3, 5639-6253\3, 8544-8902\3, 7853-8542\3, 23664-24149\3, 1-282\3, 5294-5638\3, 48919-49311\3, 2260-2703\3, 2087-2259\3, | GTR+I+G |
|  | 2 | 55769-60174\3, 10008-14323\3, 5640-6253\3, 2-282\3, 6948-7375\3, 4860-5293\3 ,2705-3614\3, 2261-2703\3, 48920-49311\3, 283-615\3, 910-2085\3, 37780-38069\3, 3615-3989\3,  3616-3989\3, 7854-8542\3, 37099-37778\3, 14652-15067\3, 617-909\3, 31238-31695\3,  49313-49914\3, 31697-32568\3, 23665-24149\3, 5295-5638\3, 43364-43956\3, 24150-26471\3, 284-615\3, 911-2085\3, 24151-26471\3 | GTR+I+G |
|  | 3 | 618-909\3, 48921-49311\3, 2706-3614\3, 14653-15067\3, 6949-7375\3, 17494-17927\3, 7378-7852\3, 285-615\3, 14326-14650\3, 43959-44316\3, 51456-51855\3, 2088-2259\3,  27885-29817\3, 23666-24149\3, 3-282\3, 37100-37778\3, 43365-43956\3, 4366-4858\3,  8545-8902\3, 22462-22768\3, 49314-49914\3 | GTR+I+G |
|  | 4 | 20631-22459\3, 912-2085\3, 49917-51453\3, 48063-48918\3, 51856-53831\3, 22771-23663\3 | GTR+I+G |
|  | 5 | 5641-6253\3, 31239-31695\3, 60177-60969\3, 60972-61425\3, 53834-54476\3, 5296-5638\3, 6256-6946\3, 4861-5293\3, 3617-3989\3, 36428-37097\3, 27096-27882\3, 2262-2703\3,  31698-32568\3, 15070-15457\3 | GTR+I+G |
|  | 6 | 17492-17927\3, 60970-61425\3, 3990-4363\3, 51454-51855\3, 53832-54476\3, 3991-4363\3, 4859-5293\3, 15069-15457\3, 43957-44316\3, 8543-8902\3, 60175-60969\3, 6254-6946\3,  4364-4858\3, 22460-22768\3, 36426-37097\3, 48061-48918\3, 29818-31236\3, 22770-23663\3, 60971-61425\3, 54478-55767\3, 38071-39353\3, 15459-17491\3, 17929-19697\3,  39355-43362\3, 32569-34896\3, 64601-65746\3 | GTR+I+G |
|  | 7 | 27094-27882\3, 15458-17491\3, 15068-15457\3, 38070-39353\3, 19700-20628\3,  19698-20628\3, 17928-19697\3, 8903-10006\3, 34898-36425\3, 17493-17927\3, 34899-36425\3, 3992-4363\3 | GTR+I+G |
|  | 8 | 15460-17491\3, 32571-34896\3, 55770-60174\3, 10009-14323\3, 24152-26471\3, 7855-8542\3 | GTR+I+G |
|  | 9 | 39356-43362\3, 54479-55767\3, 53833-54476\3, 44319-48060\3, 8904-10006\3, 26474-27093\3, 51857-53831\3, 44317-48060\3, 49916-51453\3, 61428-64600\3, 61426-64600\3 | GTR+I+G |
|  | 10 | 38072-39353\3, 29820-31236\3, 37781-38069\3, 8905-10006\3 | GTR+G |
|  | 11 | 29819-31236\3, 51858-53831\3, 37779-38069\3, 20630-22459\3, 14324-14650\3,  14325-14650\3, 48062-48918\3, 32570-34896\3 | GTR+G |
|  | 12 | 17930-19697\3, 64603-65746\3, 54477-55767\3, 26473-27093\3, 39354-43362\3 | GTR+I+G |
|  | 13 | 22769-23663\3, 49915-51453\3, 44318-48060\3, 26472-27093\3, 61427-64600\3 | GTR+I+G |
|  | **Unrooted phylogeny** | | |
|  | 1 | 10754-11941\3, 88699-90765\3, 39003-40715\3, 1-324\3, 32847-34121\3, 85138-86505\3, 85139-86505\3, 5774-6430\3, 24961-25317\3, 3052-3708\3, 107149-108537\3, 95230-95568\3, 70081-70455\3, 24706-24960\3, 21394-21699\3, 55093-55764\3, 117036-117878\3,  7253-8665\3, 82873-83265\3, 99228-103543\3, 32367-32846\3, 83653-84204\3, 76503-78686\3, 4339-4779\3 | GTR+I+G |
|  | 2 | 656-2190\3, 2192-3051\3, 64975-67551\3, 70822-72288\3, 55765-56064\3, 36168-36296\3, 70823-72288\3, 3710-4338\3, 58179-59503\3, 75300-76069\3, 15543-15956\3, 55766-56064\3, 29557-31308\3 39004-40715\3, 24962-25317\3, 10755-11941\3, 2-324\3, 32848-34121\3, 107150-108537\3, 9174-10348\3, 80403-80938\3, 88700-90765\3, 83654-84204\3,  76504-78686\3, 47340-47794\3, 5775-6430\3, 70082-70455\3, 36169-36296\3, 99229-103543\3, 4340-4779\3, 95231-95568\3, 24707-24960\3, 7254-8665\3, 28126-29201\3, 45813-46084\3, 40717-41978\3, 21701-22110\3, 21395-21699\3, 3053-3708\3, 92639-93492\3, 82874-83265\3 55094-55764\3 | GTR+I+G |
|  | 3 | 55767-56064\3, 92640-93492\3, 327-654\3, 110569-112642\3, 32369-32846\3,  116612-117035\3, 9175-10348\3, 99230-103543\3, 21396-21699\3, 35810-36167\3, 657-2190\3, 40718-41978\3, 42386-42689\3, 21702-22110\3, 95232-95568\3, 94518-95013\3,  22113-22443\3, 32849-34121\3, 39005-40715\3, 6433-7252\3, 103546-104224\3, 5773-6430\3, 16885-17230\3, 4782-5412\3, 49859-52097\3, 76505-78686\3, 4341-4779\3, 107151-108537\3, 82875-83265\3, 15959-16882\3, 3-324\3, 10351-10753\3, 113273-113687\3, 69030-69489\3 | GTR+I+G |
|  | 4 | 17232-21393\3, 104226-107148\3, 63003-64972\3, 96973-98294\3, 69029-69489\3,  14631-15541\3, 35809-36167\3, 78865-80401\3, 48466-49856\3, 25319-26941\3,  59506-63002\3, 72290-73461\3, 22445-24330\3, 78687-78863\3, 122362-123482\3,  42384-42689\3, 42385-42689\3, 16884-17230\3, 4781-5412\3, 5413-5772\3, 68747-69027\3, 32368-32846\3, 117037-117878\3, 6432-7252\3, 49858-52097\3, 86507-86928\3,  84206-85137\3, 90767-91368\3, 116611-117035\3, 37063-39002\3, 15958-16882\3,  47796-48465\3, 41980-42383\3, 94517-95013\3, 103545-104224\3, 121900-122360\3,  325-654\3, 326-654\3, 110568-112642\3, 69491-70080\3, 31310-31590\3, 78688-78863\3, 56066-57126\3, 83267-83652\3, 67553-68745\3, 91370-92637\3, 11943-14629\3,  34123-35149\3, 126453-126853\3, 5414-5772\3, 36298-37061\3, 98296-99227\3,  70457-70821\3 26943-27779\3, 95570-96972\3, 93494-94515\3, 117879-119731\3 | GTR+I+G |
|  | 5 | 29202-29555\3, 16883-17230\3, 70456-70821\3, 90766-91368\3, 36297-37061\3, 4780-5412\3, 68746-69027\3, 57128-58177\3, 121899-122360\3, 67552-68745\3, 126452-126853\3,  22444-24330\3, 26942-27779\3, 14630-15541\3, 104225-107148\3, 95569-96972\3,  37062-39002\3, 72289-73461\3, 56065-57126\3, 113271-113687\3, 34122-35149\3,  91369-92637\3, 35808-36167\3, 98295-99227\3, 93493-94515\3, 11942-14629\3,  22111-22443\3, 24332-24705\3, 6431-7252\3, 17231-21393\3, 92638-93492\3, 80402-80938\3, 63005-64972\3, 52098-53533\3, 31309-31590\3, 24331-24705\3, 94516-95013\3,  15542-15956\3, 45812-46084\3, 69028-69489\3, 15957-16882\3, 113272-113687\3,  29203-29555\3, 95015-95229\3, 96975-98294\3, 84205-85137\3, 69490-70080\3,  59505-63002\3, 83266-83652\3, 116610-117035\3, 95014-95229\3, 103544-104224\3,  110567-112642\3, 47339-47794\3, 40716-41978\3, 86506-86928\3, 21700-22110\3,  41979-42383\3, 47795-48465\3, 28125-29201\3, 49857-52097\3, 120752-121898\3,  120751-121898\3, 655-2190\3 | GTR+I+G |
|  | 6 | 31592-32366\3, 123484-124940\3, 109204-110566\3, 125961-126451\3, 54467-55092\3,  10350-10753\3, 35151-35807\3, 53534-54464\3, 113688-116609\3, 108539-109202\3,  46086-47338\3, 27781-28124\3, 8667-9172\3, 86930-88698\3, 82109-82872\3, 73462-75298\3, 119732-120749\3, 80940-82106\3, 126856-129475\3, 76072-76502\3, 76071-76502\3,  80941-82106\3, 124942-125959\3, 113690-116609\3, 124941-125959\3, 126855-129475\3, 125960-126451\3, 119733-120749\3, 123483-124940\3, 10349-10753\3, 53536-54464\3,  27780-28124\3, 35150-35807\3, 8666-9172\3, 112644-113270\3, 54466-55092\3, 2191-3051\3, 64974-67551\3 | GTR+I+G |
|  | 7 | 125962-126451\3, 2193-3051\3, 70824-72288\3, 8668-9172\3 | GTR+I+G |
|  | 8 | 3054-3708\3 | GTR+I+G |
|  | 9 | 75299-76069\3, 9173-10348\3, 3709-4338\3, 58178-59503\3, 108538-109202\3, 29556-31308\3, 46085-47338\3, 109203-110566\3, 86929-88698\3, 31591-32366\3, 73463-75298\3,  82108-82872\3 | GTR+I+G |
|  | 10 | 24708-24960\3 3711-4338\3 |  |
|  | 11 | 5415-5772\3 | GTR+I+G |
|  | 12 | 98297-99227\3, 14632-15541\3, 95571-96972\3, 7255-8665\3, 56067-57126\3, 86508-86928\3, 31311-31590\3, 55095-55764\3, 47797-48465\3, 28127-29201\3, 121901-122360\3,  93495-94515\3, 80404-80938\3 72291-73461\3, 24333-24705\3, 83655-84204\3,  90768-91368\3, 67554-68745\3, 126454-126853\3 | GTR+I+G |
|  | 13 | 84207-85137\3, 10756-11941\3, 34124-35149\3, 95016-95229\3 | GTR+I+G |
|  | 14 | 68748-69027\3, 47341-47794\3, 25320-26941\3, 122363-123482\3, 41981-42383\3,  36299-37061\3, 104227-107148\3, 63004-64972\3, 91371-92637\3 82107-82872\3,  11944-14629\3, 117038-117878\3, 96974-98294\3, 70458-70821\3 | GTR+I+G |
|  | 15 | 15544-15956\3 | GTR+I+G |
|  | 16 | 42690-45811\3, 25318-26941\3, 78864-80401\3, 52099-53533\3, 117880-119731\3,  42691-45811\3, 22112-22443\3, 48468-49856\3, 120750-121898\3, 52100-53533\3,  37064-39002\3, 122361-123482\3, 57127-58177\3, 78866-80401\3, 42692-45811\3,  59504-63002\3,17233-21393\3, 117881-119731\3, 26944-27779\3, 57129-58177\3,  48467-49856\3 | GTR+I+G |
|  | 17 | 22446-24330\3, 83268-83652\3, 88701-90765\3, 24963-25317\3 | GTR+I+G |
|  | 18 | 119734-120749\3, 27782-28124\3, 109205-110566\3 | GTR+G |
|  | 19 | 54465-55092\3, 29204-29555\3, 69492-70080\3, 36170-36296\3 | GTR+I+G |
|  | 20 | 53535-54464\3, 85140-86505\3, 35152-35807\3, 31593-32366\3, 29558-31308\3 | GTR+I+G |
|  | 21 | 45814-46084\3 | GTR+I+G |
|  | 22 | 58180-59503\3, 86931-88698\3, 46087-47338\3 | GTR+I+G |
|  | 23 | 73464-75298\3, 64973-67551\3, 112645-113270\3, 113689-116609\3, 112643-113270\3,  80939-82106\3, 126854-129475\3, 108540-109202\3, 124943-125959\3, 123485-124940\3, 76070-76502\3 | GTR+I+G |
|  | 24 | 70083-70455\3 | GTR+I+G |
|  | 25 | 75301-76069\3 | GTR+I+G |
|  | 26 | 78689-78863\3 | GTR+I+G |

**Table S7. qPCR primers to measure plasmid copy number in *Stammera* across host compartments.**

| Gene | Product | Primer | Sequence (5'-3') | Orientation | Annealing temp (˚C) | Fragment size (bp) | Target | Reference |
| --- | --- | --- | --- | --- | --- | --- | --- | --- |
| *pehA* | Endo-polygalacturonase | pg1_Chely_qpcr_F | AGCATCAAATGGACCTACATCACA | Fwd | 62 | 144 | Plasmid from *Stammera* of *Chelymorpha alternans* | This study |
|  |  | pg1_Chely_qpcr_R | ACCACTAGTTGTTCCGTTCATTGA | Rev |  |  |  |  |
| *groL* | 60 kDa chaperonin | groL1_Chely_qpcr_F | TGCTGCTTCTGTTGCTGGAT | Fwd | 62 | 124 | Chromosome from *Stammera* of *Chelymorpha alternans* | This study |
|  |  | groL1_Chely_qpcr_R | TTCCACCCATTCCTGAACCA | Rev |  |  |  |  |
| *16S rRNA* | 16S  ribosomal  RNA | f16S_StaChe | CGAGGGATGCGAGCGTTAAT | Fwd | 62,7 | 199 | Chromosome from *Stammera* of *Chelymorpha alternans* | (1) |
|  |  | r16S_StaChe | CCGCCCTTCGCCACTGATATT | Rev |  |  |  |  |

**Table S8. PCR primers to confirm the presence of the host encoded polygalacturonase in *Calyptocephala attenuata*.** Related to Figure S8.

| **Gene** | **Product** | **Primer** | **Sequence (5'-3')** | **Orientation** | **Melt temp (˚C)** | **Fragment size (bp)** | **Target** | **Reference** |
| --- | --- | --- | --- | --- | --- | --- | --- | --- |
| ***pgu1*** | Polygalacturonase | pg1_Caly_F | TGTGTAGTGTCGTCATTTCGG | Fwd | 60 | 1408 | Polygalacturonase encoded by *Calyptocephala attenuata* | This study |
|  |  | pg1_Caly_R | TTTGCCAGCGTGAGTAATGA | Rev |  |  |  |  |

**Table S9. Assembly statistics of Cassidinae transcriptomes.** Data for *Chelobasis bicolor* and *Calyptocephala attenuata* were generated in this study. Other RNAseq data sets were obtained from a previous study by (2) and Sequence Read Archive (SRA) accession numbers are indicated in the following table.

| **Beetle species** | **Total assembled bases** | **Total trinity transcripts** | **N50** | **BUSCO (%)** | **SRA accession numbers** |
| --- | --- | --- | --- | --- | --- |
| *Acromis sparsa* | 105149650 | 119737 | 1659 | 92,1 | SRR10030202 |
| *Chelymorpha alternans* | 91832612 | 65919 | 3045 | 92,6 | SRR10030205 |
| *Cistudinella foveolata* | 102002124 | 99490 | 1964 | 92,4 | SRR10030206 |
| *Discomorpha panamensis* | 66934931 | 60044 | 2037 | 88,5 | SRR10030201 |
| *Ischnocodia annulus* | 80944880 | 88954 | 1883 | 83,1 | SRR10030203 |
| *Cassida rubiginosa* | 946549465 | 1476805 | 895 | 94,8 | SRR6176960 |
| *Parachirida semiannulata* | 54754341 | 63170 | 1605 | 79,6 | SRR10030204 |
| *Chelobasis bicolor* | 100449227 | 99284 | 2076 | 93,9 | This study |
| *Calyptocephala attenuata* | 73049777 | 86241 | 1743 | 89,7 | This study |

**Table S10. Fluorescence in situ hybridization (FISH) probes used in this study.**

| **Gene** | **Product** | **Probe** | **sequence (5'-3')** | **Target** | **Dye** | **Labeling** | **Reference** |
| --- | --- | --- | --- | --- | --- | --- | --- |
| *18S rRNA* | 18S ribosomal RNA | EUK1195 | GGGCATCACAGACCTG | All eukaryotes | Atto488 | DOPE | (3) |
| *16S rRNA* | 16S ribosomal RNA | SCA600 | AAACCACCTACATGCTCTTTACGCCC | *Stammera* from 52 Cassidinae species | Cyanine 5 | DOPE | This study |
| *16S rRNA* | 16S ribosomal RNA | SAL227 | GGTCTTGAAAAAAAAAGATCCCC | Stammera from *Chelymorpha alternans* | Atto550 | DOPE | This study |

Dataset S1 (separate file). Gene clusters and their annotation representing core, accessory and singleton genes identified in *Stammera’*s pangenome.

Dataset S2 (separate file). Single copy core genes present in *Stammera*'s pangenome. Average length of the genes belonging to each gene cluster and their d*N*/d*S* values are included.

Dataset S3 (separate file). Differentially expressed symbiont genes across host compartments.

Sheet1. Differentially expressed symbiont genes when *Stammera* resides in egg caplets relative to foregut-symbiotic organs of larvae. An adjusted FDR < 0.05 and a fold-change > 0.5 were used to identify these genes. Highlighted cells in green represent differentially expressed symbiont genes also identified when comparing egg caplets relative to foregut-symbiotic organs of adults.

Sheet2. Differentially expressed symbiont genes when *Stammera* resides in egg caplets relative to foregut-symbiotic organs of adults. An adjusted FDR < 0.05 and a fold-change > 0.5 were used to identify these genes. Highlighted cells in green represent differentially expressed symbiont genes also identified when comparing egg caplets relative to foregut-symbiotic organs of larvae.

Sheet3. Differentially expressed symbiont genes when *Stammera* resides in foregut-symbiotic organs of larvae relative to adults. An adjusted FDR < 0.05 and a fold-change > 0.5 were used to identify these genes.

Sheet4. Upregulated symbiont genes when *Stammera* resides in egg caplets relative to foregut-symbiotic organs of larvae. An adjusted FDR < 0.05 and a fold-change > 0.5 were used to identify these genes. Highlighted cells in green represent upregulated symbiont genes also identified when comparing egg caplets relative to foregut-symbiotic organs of adults.

**Sheet5. Upregulated symbiont genes when *Stammera* resides in egg caplets relative to foregut-symbiotic organs of adults.** An adjusted FDR < 0.05 and a fold-change > 0.5 were used to identify these genes. Highlighted cells in green represent upregulated symbiont genes also identified when comparing egg caplets relative to foregut-symbiotic organs of larvae.

**Sheet6. Upregulated symbiont genes when *Stammera* resides in foregut-symbiotic organs of larvae relative to egg caplets.** An adjusted FDR < 0.05 and a fold-change > 0.5 were used to identify these genes. Highlighted cells in green represent upregulated symbiont genes also identified when comparing foregut-symbiotic organs of adults relative to egg caplets.

**Sheet7. Upregulated symbiont genes when *Stammera* resides in egg caplets relative to foregut-symbiotic organs of adults.** An adjusted FDR < 0.05 and a fold-change > 0.5 were used to identify these genes. Highlighted cells in green represent upregulated symbiont genes also identified when comparing egg caplets relative to foregut-symbiotic organs of larvae.

**Dataset S4 (separate file). Gene annotation of *Stammera* from *Chelymorpha alternans*.**
